# Physical modelling of multivalent interactions in the nuclear pore complex

**DOI:** 10.1101/2020.10.01.322156

**Authors:** Luke K. Davis, Anđela Šarić, Bart W. Hoogenboom, Anton Zilman

## Abstract

In the nuclear pore complex (NPC), intrinsically disordered proteins (FG Nups) along with their interactions with more globular proteins called nuclear transport receptors (NTRs) are vital to the selectivity of transport into and out of the cell nucleus. While such interactions can be modelled at different levels of coarse graining, *in-vitro* experimental data have been quantitatively described by minimal models that describe FG Nups as cohesive homogeneous polymers and NTRs as uniformly cohesive spheres, where the heterogeneous effects have been smeared out. By definition, these minimal models do not account for the explicit heterogeneities in FG Nup sequences, essentially a string of cohesive and non-cohesive polymer units, and at the NTR surface. Here, we develop computational and analytical models that do take into account such heterogeneity at a level of minimal complexity, and compare them to experimental data on single-molecule interactions between FG Nups and NTRs. Overall, we find that the heterogeneous nature of FG Nups and NTRs plays a minor role for their equilibrium binding properties, but is of significance when it comes to (un)binding kinetics. Using our models, we predict how binding equilibria and kinetics depend on the distribution of cohesive blocks in the FG Nup sequences and of the binding pockets at the NTR surface, with multivalency playing a key role. Finally, we observe that single-molecule binding kinetics has a rather minor influence on the diffusion of NTRs in polymer melts consisting of FG-Nup-like sequences.

## INTRODUCTION

The shuttling of macromolecules between the cytoplasm and nucleoplasm is controlled by nuclear pore complexes (NPCs), selective gatekeepers that permeate the nuclear envelope [1]. The NPC is a large (diameter ∼ 100 nm;mass ∼ 112 MDa in vertebrates) proteinaceous conduit that allows small (diameter ≲ 5 nm;mass ≲ 60 kDa) molecules to passively diffuse through, whilst also hindering the transport of larger molecules [2, 3]. This size-exclusion barrier arises from the dense (∼ 100-300 mg/ml) arrangement of intrinsically disordered proteins (FG Nups) that are attached to the inner NPC scaffold [4, 5]. FG Nups consist of hydrophobic (Phe-Gly) motifs that are separated by more polar/charged regions, and the relative amounts of these − hydrophobic to charged ratio − defines a cohesiveness spectrum for FG Nups, where higher amounts of hydrophobicity are associated with higher levels of cohesion [6, 7]. A Large macromolecule can travel through the NPC by binding to one or more nuclear transport receptors (NTRs), forming a cargo-complex [8]. NTRs are decorated with hydrophobic grooves that have an affinity to the hydrophobic motifs on the FG Nups, thus enabling the cargo-complex to overcome the free energy costs of entry [9, 10]. Remarkably, the NPC can facilitate the rapid transport of thousands of cargoes per second whilst also maintaining the permeability barrier. The apparent contradiction between fast transport and high selectivity is known as the “transport paradox” [3, 11].

Whilst there is a consensus regarding the importance of FG Nup-NTR interactions in NPC functionality, there is an incomplete mechanistic understanding of the exact roles these interactions play in facilitated transport. The main obstacles to this understanding are: 1) an incomplete quantification of FG Nup sequence-specific effects on the selective barrier; 2) a relative scarcity of quantitative data regarding multivalent interactions between FG Nups and NTRs; 3) a difficulty in determining the factors leading to high specificity of NTRs to the FG Nups whilst maintaining fast unbinding kinetics; 4) conflicting microscopic and macroscopic binding data on FG Nup and NTR interactions, where relatively weak per FG-motif binding (with dissociation constant *K*_*D*_ ∼ 10 mM) is observed in single-molecule studies [12, 13] as compared with strong FG Nup-NTR binding (*K*_*D*_ ≲ 10 µM) as found using other techniques [11, 14 – 18].

Physical modelling can aid in the interpretation of experiments, incorporating FG Nup and NTR heterogeneity at various levels of detail [5, 7, 15, 19 – 25]. Modelling approaches accounting for the properties of each amino acid and have reproduced elements of FG Nup morphology and FG Nup - NTR interactions, but require a calibration of many (≳ 20) interaction parameters, which poses the risk of overfitting. In addition, having a large parameter space makes it difficult to identify/explore the roles of a few key physical elements. In contrast, minimal physical modelling approaches based on a few interaction parameters, determining overall cohesion and repulsion, can help to interpret and understand current experimental observations and explore the possible outcomes of future experiments using the smallest set of governing principles.

Since both higher resolution and more coarse-grained modelling approaches aim to describe the same system, they serve complementary purposes. Here we focus on minimal-complexity models. The simplest of these models treats FG Nups as homopolymers and NTRs as uniform/isotropic particles. With experimentally determined parameter choices, these have seen surprising success in qualitatively and quantitatively reproducing binding behaviour in FG Nup and NTR in-vitro assemblies [7, 21, 22, 26]. The success of these homopolymer models suggests that many key features of NPC transport can be described by mesoscopic polymer-colloid physics, and naturally raises the question of the roles that FG Nup and NTR heterogeneity have on NPC functionality.

Here we present modelling approaches to explore the roles of FG Nup sequence patterning, surface heterogeneity (or “patchiness”) of NTRs, and multivalency in regulating FG Nup-NTR binding and associated binding kinetics, and their effects on the equilibrium diffusion of an NTR in an FG polymer melt. The structure of the paper is as follows: first we present the methodology of our approach, second we set the parameters and validate the model through comparison with experimental data, third we provide analytical frameworks for interpreting the results, fourth we use the model to explore a more physiological FG sequence binding to NTRs, and finally we make predictions of equilibrium diffusion of an NTR in an FG polymer melt and compare with experimental data.

## METHODS

### Coarse-grained model

Intrinsically disordered proteins that contain hydrophobic FSFG repeats that are separated by neutral regions containing other amino acids [13] are modelled as freely-jointed heteropolymers (see Figure 1). To minimize the complexity of the model, we only include two types of beads − cohesive and non-cohesive - and impose that all polymer beads have the same size, as is commonly done in homopolymer models [7,21,22]. To examine the effects of a particular choice of the monomer size, *d*, we test sizes ranging from the approximate diameter of an amino acid (one amino-acid-per-bead (1apb), *d* = 0.38 nm) to 2 amino acids (2apb, *d* = 0.76 nm) and finally to that of 4 amino acids (4apb, *d* = 1.02 nm, *i,e* in between the size of 4 close packed amino acids and a straight line of 4 amino acids). Two neighbouring polymer beads are connected by a stiff harmonic spring defined by the potential 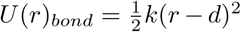, where *k* is the bond strength and *r* is the distance between two neighbouring monomers.

**FIG. 1.**
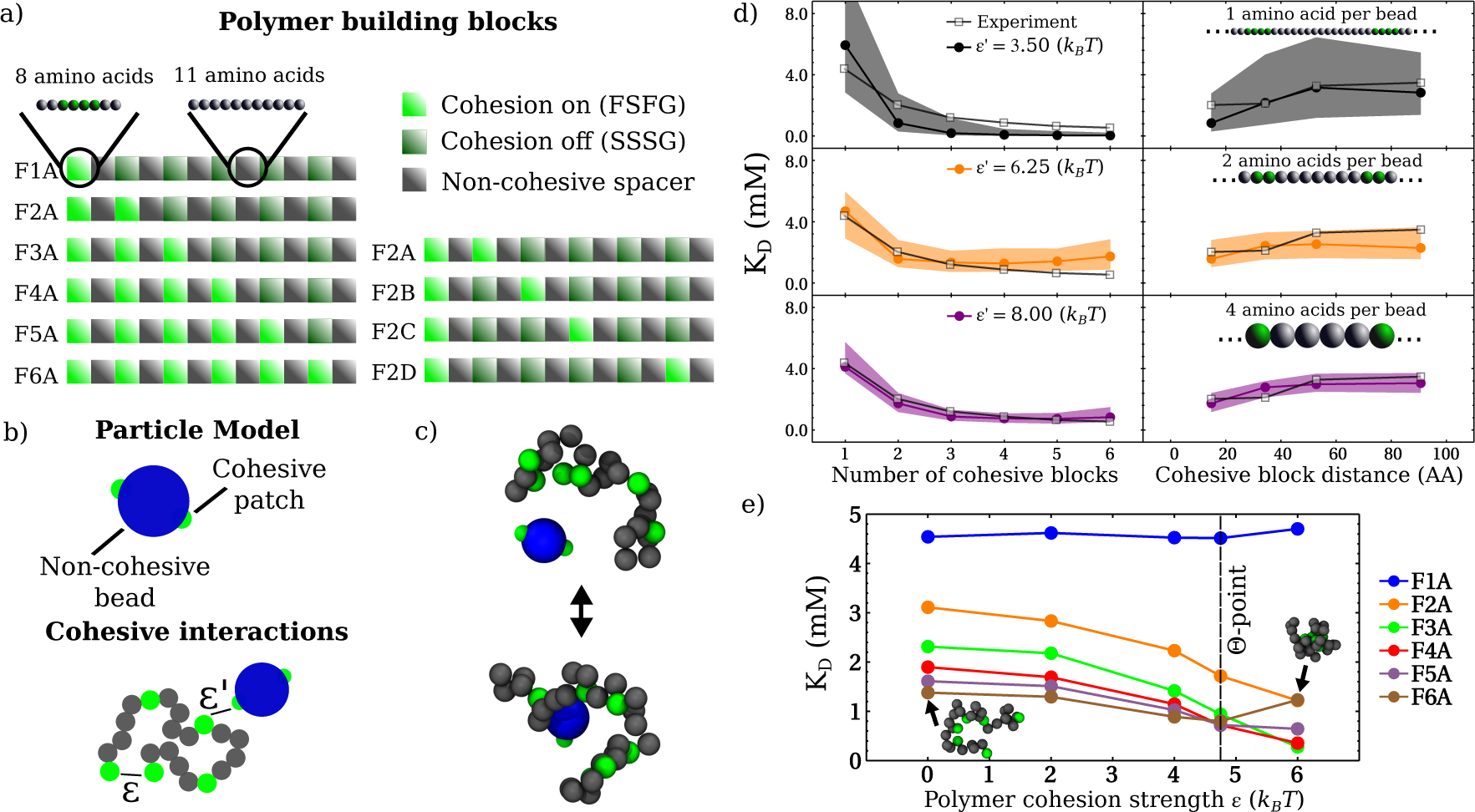
Set up of the physical model used in this work. a) Polymer sequences comprised of three types of (amino-acid containing) blocks that mimic synthetic protein sequences for which experimental data are available [13]. b) NTF2 is modelled as a patchy particle and the polymer sequences as chains with cohesive and non-cohesive beads. *ϵ* and *ϵ*′ define the attraction strength between cohesive beads with the chain, and between these cohesive beads and the cohesive patches on NTF2, respectively. c) MD snapshots of unbound and bound NTF2 with an F6A polymer (4 amino acids per bead). d) Parameterization of the particle-polymer cohesion strength *ϵ*′, shown for different choices of polymer bead size, using NMR data on the dependence of *K*_*D*_ on the number of cohesive blocks (left) and on the separation of two blocks along the sequence (right) [13]. Shaded bands denote *ϵ*′ *±* 0.25*k*_*B*_ *T*.) *K*_*D*_ as a function of *ϵ*, based on simulations with 4 amino acids per bead, with a fixed *ϵ*′ as determined in d. The Θ-point (vertical dashed line) and inset snapshots refer to the F6A sequence.

We incorporate the size and binder coverage of a nuclear transport receptor in a similarly minimal fashion. In this model, a nuclear transport receptor is treated as a rigid body composed of a sphere of diameter 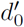 that has *N* spheres (“patches”) of diameter *d*′ = 0.38 nm (where 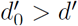) fixed at specified points on its surface.

We imposed excluded volume interactions between all constituent particles via the Weeks-Chandler-Anderson potential

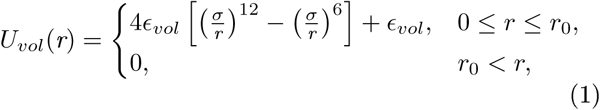

where *ϵ*_*vol*_ = 250 *k*_*B*_*T* is the interaction strength, 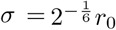, and *r*_0_ is the distance at which the excluded volume interaction acts. The addition of *ϵ*_*vol*_ to the potential ensures that *U*_*vol*_(*r* = *r*_0_) = 0.0 *k*_*B*_*T*. We also imposed a cohesive interaction of strength *ϵ* between cohesive polymer beads and an additional cohesive interaction of strength *ϵ*′ between cohesive polymer beads and cohesive patches on the particle through the Morse potential

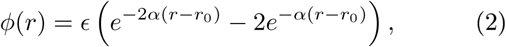

which is truncated and shifted to ensure continuity in both the potential and force for *r* ≤ 2*r*_0_, where 2*r*_0_ is the cut-off of the cohesive interaction. The resulting cohesive pair potential is then given by

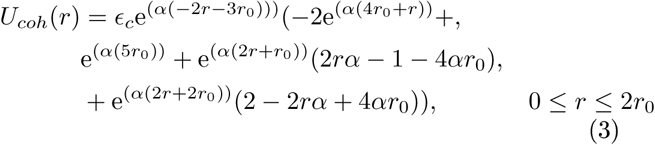

where *α* = 6.0 nm^−1^ is the decay of the Morse potential and *ϵ*_*c*_ = {*Aϵ*, *A*′*ϵ*′}, in which *A* and *A*′ are constants, recorrects the minimum of the potential, that is initially set through either of the two interaction parameters {*ϵ*, *ϵ*′}, due to the truncation and shifting procedure. Combining the pair potentials for excluded volume *U*_*vol*_ and cohesion *U*_*coh*_ then leads to the overall interaction given by

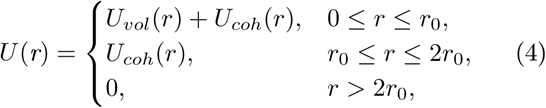

of which an example plot is shown in the supplementary information (SI) Figure 1. Relevant length scales for the different polymer and particle models are given in SI Table 1.

### Simulation details

MD simulations were performed using the LAMMPS package [27]. We subjected the patterned polymer and patchy-particle system to dynamics at a constant temperature, *T*, through the implementation of the NVE (constant number of particles N, constant colume V, constant total energy E) time integration algorithm with a Langevin thermostat. This combination results in the total force **F** acting on a particle as given by

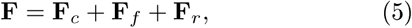

where **F**_*c*_ is the conservative force due to the inter-particle pair potentials, **F**_*f*_ = −(*m/γ*)***v*** is the frictional drag force, and 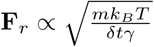 is the force due to solvent particles at a temperature *T* randomly colliding with the particle. Simulations were performed with dimensionless parameters with *T* = 1 and *γ =* 1, where *γ* is the friction coefficient, and a simulation timestep of *δt* = 0.002*τ*_0_, where *τ*_0_ = 1.707 × 10^−9^ s is the unit of time as defined in the supplementary information of our previous work [7]. The patchy-particle was treated as a rigid body so that the resultant force and torque of the body is the sum of the forces and torques of the constituent particles.

To simulate a single polymer (of total volume *v*) and a single particle (of total volume *v*′) we initially placed them in a box, with periodic boundary conditions, of size *L*^3^ where typically *L*^3^ *> C*(*v* + *v*′), with *C >* 80 (corresponding to *L >* 20 nm). Next we performed a simulation run for 5 × 10^6^ timesteps to equilibrate the system, which was checked upon inspection of the total energy, a further 30 × 10^6^ timesteps were used for data analysis.

To simulate a bulk system consisting of *N*_*p*_ polymers, and *M*_*p*_ particles both at packing fractions *η* in a periodic box of length *L* = 40 nm, we first placed *N*_*p*_ = *ηL*^3^*/V*_pol-beads_ polymers and *M*_*p*_ = *ηL*^3^*/V*_pol-beads_ in a cubic box of length *L*_0_ = 120 nm (centered about the origin), where *V*_pol-beads_ is the sum of constituent bead volumes of the patterned polymers and *V*_pol-beads_ is the sum of constituent bead volumes of the patchy-particles. Then, in order to shrink the box to a volume of *L*^3^ to obtain a relatively high density (and the desired packing *η*), we performed an initial run which applied an external force pushing all beads to the centre of the box whilst avoiding particle overlaps. After this we performed a simulation reducing the box length from *L*_0_ to *L* and let the system equilibrate (20 × 10^6^ timesteps), checked upon inspection of the total energy.

## RESULTS AND DISCUSSION

### Minimal-complexity models account for NTF2 binding to short FSFG containing sequences

To develop and test minimal-complexity models for how NTRs bind to FG domains, we referred to experiments on the 1:1 binding of NTRs – specifically of NTF2 - to short (≈120 amino acids) FSFG-containing sequences [13]. These sequences contained various patternings of cohesive (sticker) blocks consisting of cohesive FSFG with 2 flanking −non−cohesive−amino acids on either side, and non-cohesive spacer blocks consisting of 11 non-cohesive amino acids. In addition, each FSFG block could be substituted with a non-cohesive SSSG alternative. The different sequences are schematically depicted in Figure 1a. Briefly, in these sequences, the amount and relative locations of the cohesive blocks were systematically varied; NMR was used to measure the respective *K*_*D*_s for NTF2 binding to the various sequences (with additional validation by calorimetry) [13].

We model these sequences as beads-on-a-chain with different levels of coarse-graining (1, 2, and 4 amino acids per bead), where the sequence heterogeneity is incorporated at minimal complexity using appropriate alternations of cohesive (for FSFG-containing blocks) and non-cohesive (for the other blocks) beads. Given that NTF2 has at least two FxFG binding sites [28], we treat NTF2 as a sphere – of diameter 3 nm – with two cohesive patches. The intra- and intermolecular affinities in this system are defined, respectively, by the strength of interactions between the cohesive beads on the chain, *ϵ*, and by the strength of interactions between the cohesive beads on the chain and the cohesive patches on the NTR, *ϵ*′, see Figure 1b.

In accordance with previous findings that native and synthetic FG Nup domains behave similarly to Θ-point homopolymers [7, 21, 22, 25, 29, 30], where intramolecular repulsion balances intramolecular attraction, we set the polymer-polymer cohesion strength *ϵ*= 4.75 *k*_*B*_*T* such that the F6A chain is at the Θ-point (see SI Figure 1). We chose to set *ϵ* based upon the sequence with six cohesive blocks (F6A) as it more closely resembles the symmetry of a homopolymer chain as compared to the other sequences. This leaves us with a single free parameter, the polymer-particle attraction strength *ϵ*′, to adjust dissociation constants *K*_*D*_ from MD simulations to those derived from experiment (see Figure 1c and d). In MD, *K*_*D*_ is essentially computed from counting the number of unbound states to bound states that occur in a single, long, simulation trajectory [7, 31,32] (see SI for details). Encouragingly, such one-parameter fits are sufficient for the simulations to quantitatively reproduce the behaviour of *K*_*D*_ as a function (i) of the number of FSFG motifs in the sequence and (ii) of the separation of two FSFG motifs along the chain. Moreover, this agreement was robust against the level of coarse-graining of the protein sequences, with concomitant adjustments of *ϵ* and *ϵ*′ that were parametrised by the Θ-point for F6A and by the best match with experimental *K*_*D*_ curves, respectively. We note that binding affinities (*K*_*D*_) exhibited greater sensitivity to changes in *ϵ*′ for higher resolution (less coarse-graining) polymer models (see Figure 1d). For computational convenience, we focus on the coarsest model used here, a polymer with 4 amino acids per bead (4apb).

For this model, we also investigated how NTF2 binding depends on the intra-polymer cohesion parameter *ϵ*. For the sequence with a single cohesive block (F1A), the choice of *ϵ* should not – and does not – affect *K*_*D*_. For sequences containing ≥ 2 cohesive blocks, *K*_*D*_ decreases (hence binding affinity increases) upon increasing *ϵ* (see Figure 1e). This may be attributed to the higher local concentration of particle binding sites as the polymer assumes more compact conformations. For the case of F6A, however, *K*_*D*_ increases (hence binding affinity decreases) when is increased beyond the Θ-point (*ϵ*≥4.75 *k*_*B*_*T*). This increase in *K*_*D*_ can be explained by the preference for cohesive beads on the polymer to remain bound to one another, gradually forming a tightly bound globule with the non-cohesive polymer regions covering it, thus reducing the accessibility of available binding sites for NTF2 (see SI Figure 1 and Figure 1e). Accordingly, for such high intra-polymer cohesion, the *K*_*D*_s also go up with an increasing number of cohesive blocks for F3A –F6A, inverting the trend observed below the Θ-point (Figure 1e). Overall, this decrease in polymer-particle affinity will be defined by the increased polymer cohesion and by the entropic cost of confining non-cohesive regions [33, 34].

Having thus established a coarse-grained model that accounts for sequence-dependent variations of how NTF2 binds to FSFG-containing sequences, we next set out to investigate the importance of surface heterogeneity (for NTF2) and sequence heterogeneity (for the FSFG-containing sequences) to quantitatively account for the experimental data. Firstly, we varied the proximity of the two cohesive patches on the NTF2 -mimicking particle, through the angle *θ*. We observed little change in the resulting *K*_*D*_ until the patches were very close, *i*.*e*., for *θ ⪅π/*5 (≡36°) (see Figure 2a). For *θ ⪅ π/*5, the binding became so strong that NTF2 was hardly released from the polymer sequences in our simulations; this strong binding is attributed to the increase in binding energy when a polymer block binds to two NTF2 patches simultaneously. This strong binding was also articulated in the reduction in polymer size (the radius of gyration *R*_*G*_) for sequences with increasing amounts of cohesive blocks in the sequence. For larger values of *θ*, with weaker binding, there was an initial mild shrinkage with the number of cohesive blocks, followed by a more pronounced expansion (size increase *<* 10%). Such expansion of polymer size upon binding has also been observed in experiments on NTF2 binding to similar FG domains [35], providing further support for our model.

**FIG. 2.**
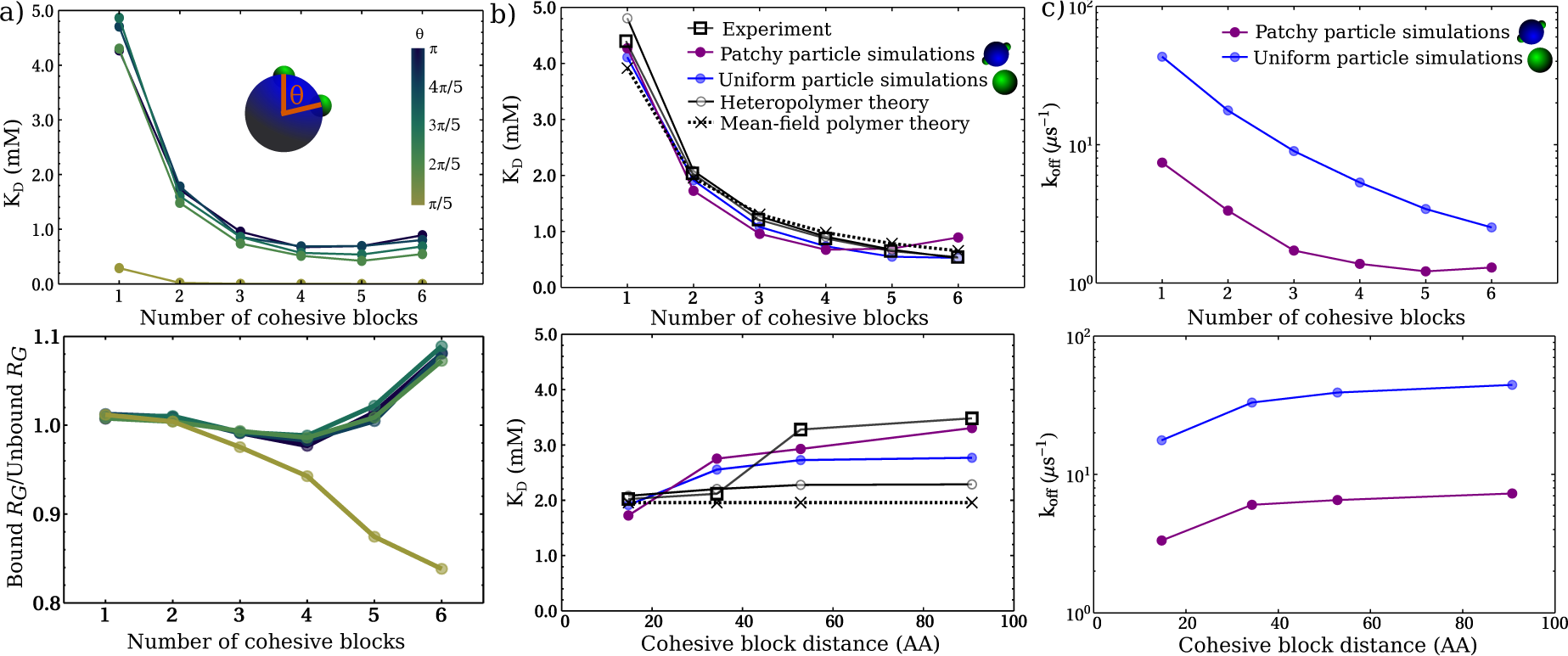
Effects of surface heterogeniety on a nuclear transport receptor. a) Effects of varying the patch-patch proximity on binding affinity and polymer size (as measured by the radius of gyration *R*_*G*_). b) Effects of surface heterogeneity on the particle and sequence heterogeneity in the polymer using simulations and polymer analytical models. For the patchy particle simulations *θ* = *π* rads (≡ 180°) is used. In the uniform particle cases, the particle interacts with the cohesive polymer beads through its center of mass. As is shown, the cohesion strengths in all cases are set to best match experiments [13]. c) Comparing the kinetics between the best match (of the experimental data) uniform particle and patchy particle model used in simulations. The off rate was calculated as *k*_off_ = ⟨*t*_res_ ⟩^−1^, where ⟨*t*_res_ ⟩ is the ensemble averaged interaction residence time.

Noting that we did not observe a major change in *K*_*D*_ with minor variations of the position of the patches on NTF2 (for *θ > π/*5), we also attempted to model the experimental data ignoring surface heterogeneity altogether and instead modelling NTF2 as a homogeneous/uniform cohesive sphere (of the same size as the non-cohesive bead in the patchy-particle above). In the case of the uniform particle model of NTF2, the cohesive interaction – as imposed through a pair potential – is between the centre of the large bead (≈8 times larger than the cohesive NTF2 patch) and the centre of the cohesive polymer bead, with an interaction cutoff distance that is ≈3-fold larger than the interaction cutoff distance in the patchy-particle model. As before, we adjusted *ϵ*′ to best reproduce the variation of *K*_*D*_ with the number of FSFG blocks and with their separation along the sequence, resulting in a value of *ϵ*′= 5.0 *k*_*B*_*T* for the uniform NTF2 model (Figure 2b). These results suggest that surface heterogeneity *per se* is not a relevant factor to determine binding of NTRs to short FG domains, as long as the overall affinity of the NTRs to the FG domains is in the appropriate range. This is largely consistent with experimental observations, which indicate that there are multiple ways to tune NTR surface properties to achieve similar transport properties [36].

### Analytical models quantitatively describe effects of multivalency

To explore how the heterogeneity in the polymer sequence dictates the binding affinities as observed in the experimental data, we used an analytical theory that enables quantification of sequence-specific effects on binding [37, 38] (see SI for mathematical details). In line with our results above, we considered the NTF2 as a uniform particle (one bead) binding to a patterned polymer defined by a patterning of cohesive and non-cohesive beads. Intra- and intermolecular interactions were incorporated in an approximation of the second virial coefficient *B*_2_ via two terms. The first term is based on a mean-field contribution, where the heteropolymer is approximated as a smearing of an uncorrelated set of *N*_coh_ cohesive monomers that has an effective affinity of strength ∝ *ϵ*′ *N*_coh_ and a range determined by the pair potential (see SI Table 1), for the NTF2 particle only. The second term is sequence-specific and is based on second-order correlations arising between the polymer monomers and also between the polymer monomers and the NTF2 particle. Therefore the intramolecular polymer correlations implicitly depend on the intra-polymer cohesion strength and on an excluded volume parameter *ω* accounting for the finite size of the polymer beads; and the correlations between the NTF2 particle and cohesive monomers of the polymer depend on *ϵ*′. In its simplest form, *B*_2_ is given as (as explicitly shown in the SI)

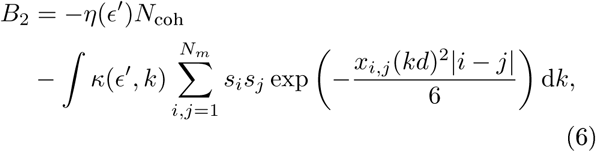

where *s*_*i*_ encodes whether a polymer bead *i* is cohesive *s*_*i*_ = 1 or non-cohesive *s*_*i*_ = 0 (with 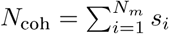),*N*_*m*_ is the total chain length (*N*_coh_ + *N* _non-cohensive_), *d* is the bond length, *x* is a renormalization factor of the bond length that corrects the Gaussian (ideal polymer) correlations for intramolecular excluded volume (dependent on the excluded volume parameter *ω*) and intramolecular cohesion (dependent on *ϵ*) [38], and the integral is over the wave number *k* defined as the inverse monomer-monomer distance. The functions *η* and *κ* are explicitly given in the supplementary information. To calculate the dissociation constant between one polymer and one particle, we use (details in the SI)

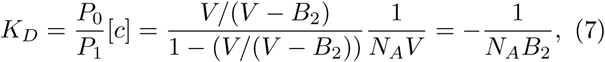

where *P*_1_ is the binding probability, *P*_0_ = 1 − *P*_1_ is the unbinding probability, [*c*] = 1*/N*_*A*_*V* is the concentration (with units mol ·nm^−3^), *N*_*A*_ is Avogadro’s number, and *V* is the system volume [31, 32, 37, 39]. Note that, since we are considering the binding of a single polymer to a single particle in a relatively large volume, we use the ratio of the unbinding to binding probabilities *K*_*D*_ ∝ *P*_0_*/P*_1_ (Boltzmann approach), which is practically the same as using 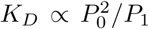 (law of mass action approach) when in the dilute limit (where 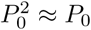) [31, 32]. The last equality in equation 7, which states that 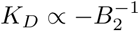, is also valid in the dilute limit [32, 37, 39]. In order to compare with the experimental *K*_*D*_ data (Figure 1d), equation 7 is converted to molar units (M) using 1 nm^−3^ ≡ 10^−24^ liters.

The intra-chain interaction parameters, {**ϵ*, ω*} can be estimated based on a matching of *R*_*G*_ between the heteropolymer theory with those calculated for the simulated sequences (as was similarly done in [38]); and the polymer-particle cohesion strength **ϵ**′ is left as a fitting parameter to describe *K*_*D*_ as a function of the number of cohesive blocks. With such parametrization, this analytical heteropolymer model accurately describes the experimental data as a function of the number of cohesive blocks (Figure 2b, top). For comparison, we next recalculated *K*_*D*_ without the second term in equation 6, with no dependence on, {**ϵ*, ω*}, and thus approximating the heteropolymer as a smearing of a set of uncorrelated monomers. Remarkably, this much simplified model reproduces the experimental data equally well as the heteropolymer model (Figure 2b, top). As per equations 6-7, the latter agreement implies that we can describe the increase in polymer-particle affinity with increasing numbers of cohesive blocks in the polymer using a simple approximate relation *K*_*D*_ ∝ 1*/N*_coh_. Indeed, the binding trends as seen in experiments can be well-fitted to a function *K*_*D*_(*N*_coh_) = *A/N*_coh_, with *A* being a fitting constant (see SI Figure 2).

Understandably, the mean-field theory (based on the first term in equation 6) could not reproduce the trends in *K*_*D*_ with the distance between (a constant number *N*_coh_ = 2 of) cohesive blocks being varied along the sequence. However, the heteropolymer theory - including second-order correlations only − correctly predicts an increase of *K*_*D*_, though of smaller magnitude than the experimental (and MD) data (Figure 2b, bottom). This provides a physical explanation of why binding affinity increases when the FSFG motifs are bought closer together on the sequence, the spatial correlations between the motifs are higher which results in an increase in local concentration of motifs around the NTR [13].

### Effect of NTR surface heterogeneity on unbinding kinetics

**A**lthough the most simplified models above accurately describe binding equilibria, it remains unclear to what extent NTR surface heterogeneity needs to be incorporated in a model to account for (un)binding kinetics. To address this question, we calculated the off-rate *k*_off_ = ⟨*τ*_res_ ⟩ ^−1^ from MD simulations, where *τ* is the ensemble-averaged time over which the binding patches of the NTR remained in the interaction range of at least one cohesive bead on the patterned polymer (Figure 2c). This results in predicted residence times of ⟨*τ*_res_⟩ *<* 1 μ*s*. With *K*_*D*_ 1 m as above, such residence times imply on-rates *k*_on_ ∼ 1 nM ^−1^s^−1^ that are consistent with estimates based on stopped-flow experiments [12]. We observe that the off-rates for the uniform NTF2 -equivalent particle (for *N*_coh_ ≤ 3) are an order of magnitude faster than that for the NTF2 with two separate binding patches, and are at least three times greater (for *N*_coh_ *>* 3), despite both particles having calculated *K*_*D*_ values that match the same experimental data (see Figure 2b). This implies that the more the NTR-FSFG affinity is smeared out accross the NTR surface, the easier it is to unbind the NTR from the FG motifs for a given equilibrium binding constant. In other words, by relying on more spread-out and weaker interactions, FG motifs can achieve the same thermodynamic selectivity for NTRs as would result from having fewer but stronger interactions, yet allow for faster off-rates. This provides at least part of the explanation for the “transport paradox” [3, 11], where many but weak binding spots allow FG motifs in the NPC to have high selectivity for NTRs, yet still facilitate fast off-rates and hence fast transport.

### Accounting for more complex FG sequences and NTRs

**H**aving established our coarse-grained models by comparison with experimental data on short FSFG-containing sequences and NTF2, we next carried out according simulations for a more complex FG sequence and more complex NTRs (Figure 3a). Specifically, we devised a longer (≈720 amino acid) sequence with a distribution of 18 FSFG cohesive blocks that is consistent with native Nsp1, an FG Nup that recapitulates salient features of the NPC selective barrier in artificial NPC mimics, [26 40, 41]. For NTRs, we chose NTF2 as well as Karyopherin-95 (Kap95, or Importin-*β*), an important and extensively studied cargo-carrying NTR. We model Kap95 as a patchy-particle (size ≈5 nm) with *N* cohesive patches embedded on its surface, with *N* ranging between 4 and 10 as is expected for Kap95 based on structural data and simulations, [9, 10, 42] (Figure 3a). To further probe the effects of surface heterogeneity, and of the proximity of cohesive patches, we used two Kap95 models one with the *N* patches equidistant from one another over the whole surface (Kap) and another with the same number of *N* patches spread equidistantly on one half of its surface (Kap_half).

**FIG. 3.**
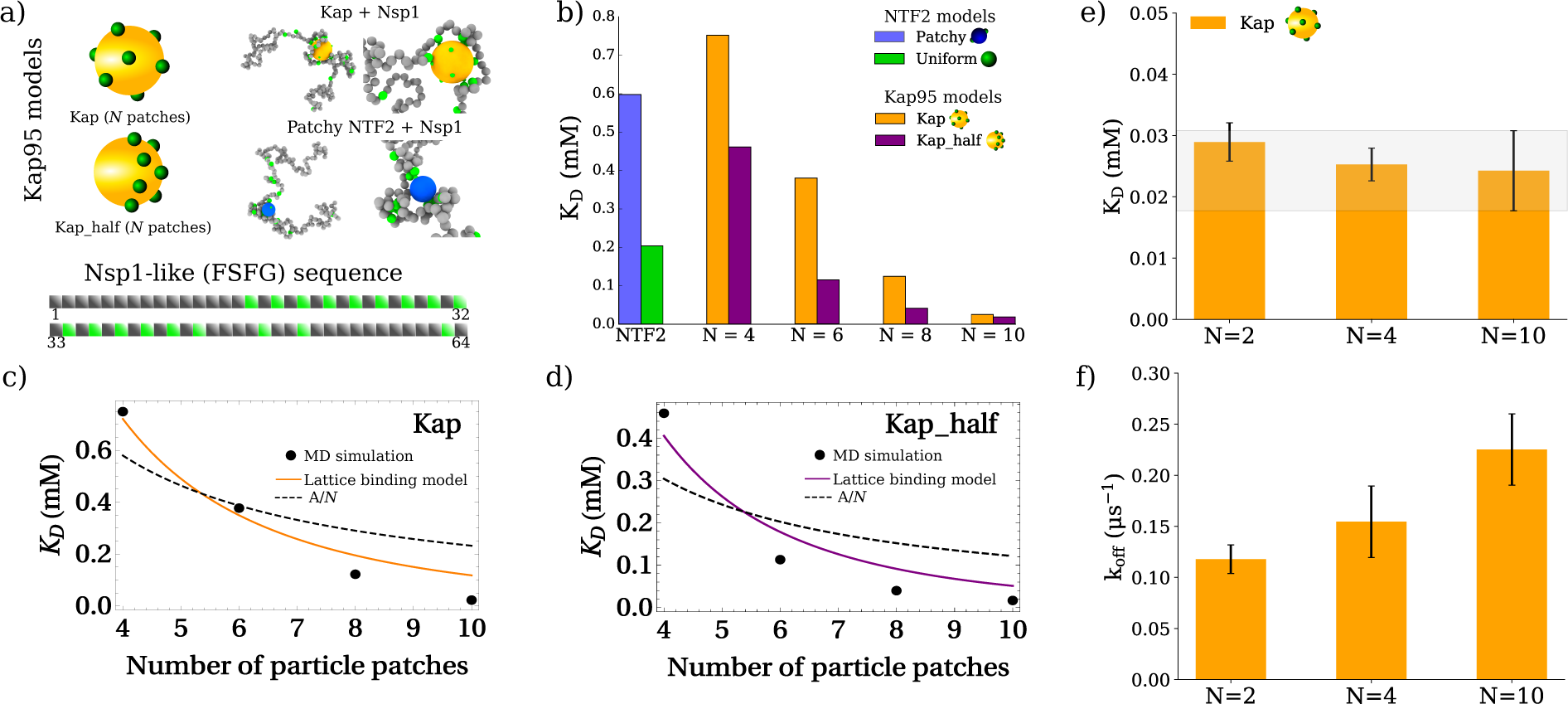
Single-molecule binding predictions for an Nsp1-like FSFG-sequence and NTR particles with varying surface properties, based on coarse-grained MD simulations. a) Illustration of the particle models for NTRs and the Nsp1-like sequence. Kap95 is modelled as a particle with *N* binding patches uniformly distributed on its surface (Kap) or a particle with *N* patches uniformly distributed on one half of its surface (Kap_half). The Nsp1-like sequence is built from the polymer building blocks shown in Figure 1a (with *N* _coh_ = 18 cohesive blocks). (Inset) MD snapshots of Nsp1 binding with selected NTR models. b) *K*_*D*_ predictions for Nsp1 and the NTR models, with **ϵ** = 4.75 *k*_*B*_ *T* for polymer − FSFG-FSFG − cohesion and **ϵ**′ = 8.0 *k*_*B*_ *T* for polymer-particle cohesion (with **ϵ**′ = 5.0 *k*_*B*_ *T* for the uniform NTF2 particle so as to match the *K*_*D*_ s of the NTF2 patchy particle with the short sequences (Figure 2b)). c,d) Fitting the *K*_*D*_ data from MD simulations (b) with the analytical lattice binding model, where the *N* _coh_ = 18 polymer cohesive blocks and cohesive particle patches interact within a sub-volume of a lattice. e,f) When **ϵ**′ is adjusted to yield the same, within standard deviation (grey band), *K*_*D*_ s (as determined by MD simulations) for *N* = {2, 4} as *N* = 10 in the Kap model (e), the off-rate increases with *N* (f). In (e,f) **ϵ**′ = 10.45, 9.55, and 8.0 *k*_*B*_ *T* for *N* = 2, 4, and 10 patches respectively.

For simplicity, we assume a rather generic nature of the interactions between FSFGs and between FSFG and binding patches on NTRs, hence we use the same = 4.75 *k*_*B*_*T* and **ϵ**′= 8.0 *k*_*B*_*T* as used for the coarsest model of the short FSFG sequences and patchy NTF2 (see Figure 1), and = 5.0 *k*_*B*_*T* for the uniform NTF2 particle as before. Firstly, considering the 2-patch NTF2 particle (with *θ* = *π* rads), we do not observe a significant drop (of ≈ 0.2 m difference) in *K*_*D*_ going from the F (with *N*_coh_ = 6) to the Nsp1-like sequence (with *N*_coh_ = 18) (see Figure 3b), whereas with a scaling *K*_*D*_ ∝ 1*/N*_coh_, we would expect a 3-fold drop in *K*_*D*_. This observation could be due to the patchy NTF2 (2 patches) not being able to make as efficient use of cohesive blocks when the latter become abundantly available, *i*.*e*, the binding patches on NTF2 start to saturate. Reassuringly, this observation is supported by experiment where an equivalent F12A sequence (12 blocks) exhibited a *K*_*D*_ that was similar in value to the F6A sequence [13]. It is worthwhile to note that whilst the Nsp1-like sequence contains more cohesive blocks, it is also 6 times longer than the F1A-F6A sequences and contains a few larger non-cohesive spacer regions that might contribute to some reduction in binding affinity, due to entropic effects. Such saturation is much less of a limitation on a uniformly cohesive NTF2 equivalent, which indeed shows an ≈ 3-fold increase in affinity to the Nsp1-like sequence as compared with the F6A sequence, consistent with the scaling *K*_*D*_ ∝ 1*/N*_coh_. This indicates the limitation of the uniform-NTR approximation it overestimates the binding of NTF2 to long FG domains when there is a large (≱10) excess of FSFG motifs that can access the NTR.

For the Kap95 models, using a constant **ϵ**′= 8.0 *k*_*B*_*T* as noted above, we observe a decrease in *K*_*D*_ (increase in overall binding energy) upon increasing in *N*. Apart from depending on the number of binding sites *N* on an NTR, *K*_*D*_ also depends on the distribution of these sites over the surface for the Kap95 model particles. Such dependence is rather pronounced when comparing the Kap and Kap_half models. As observed when bringing the binding sites of NTF2 in close proximity (Figure 2a), we observed a significantly stronger binding when all *N* binding sites are located on one side of the NTR (Kap_half) compared with the case where these binding sites are distributed uniformly over the whole surface (Figure 3b) [42]. For the Kap and Kap_half particle with *N* = 4 patches the *K*_*D*_ is comparable to that of patchy NTF2, despite having a twofold increase in patch numbers. This observation is the result of the difference in sizes between Kap95 and NTF2 particles, which provides the rationale as to why larger NTRs require more cohesive binding spots [10].

Encouraged by the previous success of our analytical models, we attempted to describe the Kap and Kap_half data using the inverse relationship (*K*_*D*_ ∝ 1*/N*) that would result from a mean-field treatment of the particle patches (see first term of *B*_2_ in equation 6). We found that the simple fitting function *K*_*D*_(*A, N*) = *A/N* (*A* being a fitting constant), that well described the experimental and simulation trends of *K*_*D*_ on the number of cohesive polymer blocks (Figure 2b and SI Figure 2), resulted in a poor fit to the Kap95 simulation data (see Figure 3(c,d)).

We therefore attempted to develop an analytical model that could describe both the *K*_*D*_ dependence on the number of cohesive polymer blocks *N*_coh_, as seen in the polymer data, and on the number of cohesive patches on the NTR *N*, as seen in the Kap95 data (see the supplementary information for a complete derivation). In this model one considers a system volume *V* where the *N*_coh_ cohesive units in the polymer and the *N* cohesive units on the particle each occupy a sub-volume *V*′, which is assumed − for simplicity − to be the same for both (see SI Figure 3). Again, for simplicity, the volume *V* is discretized into *M* small cubes of volume *v* _*vo*x_ whose centers form a cubic lattice. In the model, henceforth called the lattice binding model, the polymer cohesive units are assumed uncorrelated and can occupy any of the *M*′ = (*MV*′)*/V* sites within *V*′; the particle units are assumed fixed relative to one another (analogous to the ap model in simulations) and are symmetrically distributed amongst the *M*′ sites. The polymer and particle units can only interact with one another when their respective sub-volumes overlap, when an individual polymer unit occupies the same lattice site of a particle unit it is considered to be bound. Hence, the partition function for this system is given by

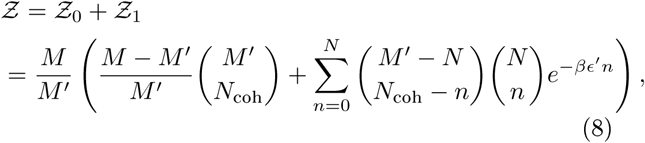

where 𝒵_0_ is the number of non-interacting microstates, and 𝒵_1_ is the number of interacting microstates. The first factor *M/M*′ is the number of ways of placing the polymer sub-lattice (*M*′) in the system lattice (*M*); the second factor (*M − M*′)*/M* is the number of ways of placing the particle sub-lattice (*M*) in the system lattice without over-lapping with the polymer sub-lattice (hence *M* − *M*′ rather than *M*); the first binomial coefficient is the number of arrangements of the *N*_coh_ units in the polymer sub-lattice; the second binomial coefficient is the number of arrangements of the *N*_coh_ − *n* unbound polymer units amongst the *M*′ − *N* free sites; the third binomial coefficient is the number of arrangements of the *n* bound polymer units amongst the *N* particle units; the last factor is the Boltzmann weight for the system energy *E* = **ϵ**′*n* (in units of *k*_*B*_*T*), where **ϵ**′is the negative binding energy. The probability that at least one polymer unit is bound is

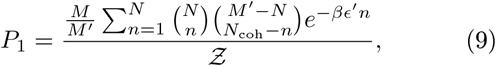

and the probability for no polymer units being bound is simply *P*_0_ = 1 − *P*_1_. Using the relation *K*_*D*_ = (*N*_*A*_*V*)^−1^*P*_0_*/P*_1_ (as in equation 7) one obtains, for arbitrary *N*_coh_ and *N*, an equation for *K*_*D*_ given as (details in the supplementary information)

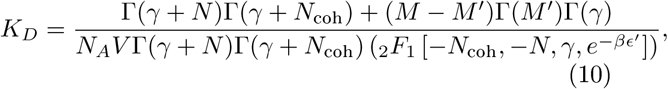

where Γ(*x*) is the Gamma function, _2_*F*_1_[*a, b*; *c*; *d*] is the Gauss hypergeometric function, and *γ* = 1 + *M*′ − *N* – *N*_coh_A s a first test for consistency, we checked whether equation 10 reproduced the *K*_*D*_ ∝ 1*/N*_coh_ relationship as shown before. Indeed, by setting *N* = 1 in equation 10 we obtain

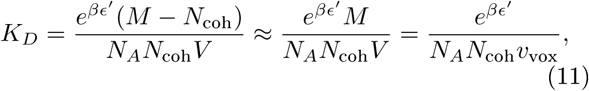

where the approximation *M >> N*_coh_ is valid in the dilute limit. Next, we performed a one parameter fit of equation 10 to the Kap and Kap_half simulation data, by setting *N*_coh_ = 18 (for the Nsp1-like sequence), and found that it reasonably describes the trends observed in the coarse-grained MD simulations (Figures 3c,d). We also checked that the results from the lattice binding model were robust against changes to the simplified assumption of how the polymer and particle units bind, *i*.*e*, by accounting for a microscopic probability of the units becoming bound (see supplementary information). Comparing the data in Figure 3b with other experiments, we note that the *N* = 10 result (*K*_*D*_ = 24 μ M) is in reasonably good agreement with experimental affinities measured for FG Nups and importins *K*_*D*_ *⪆* 4 μ M [11, 14].

Next, considering the unbinding kinetics using the MD simulations as for the case for NTF2 (Figure 2c), we find that for NTRs with more spread out binding sites the unbinding off-rates *k* are higher when comparing NTR models that yield similar *K*_*D*_s. That is, if the respective **ϵ**′s are increased for the *N* = 2 (**ϵ**′ = 10.45 *k*_*B*_*T*) and *N* = 4 (= 9.55 *k*_*B*_*T*) Kaps in order for them to have the same *K*_*D*_ as the *N* = 10 (**ϵ**′= 8.0 *k*_*B*_*T*) particle (Figure 3e), the *k* increases with *N* (Figure 3f). This confirms our result (see also Figure 2c) that to facilitate fast kinetics in spite of having strong binding, it is beneficial for NTRs to have many weak binding sites on their surface rather than a few stronger ones.

### Diffusion of NTRs in a polymer melt

Following the success of our models in replicating experimental data on single NTRs binding to single FG domains, and having used this to study binding equilibria and unbinding kinetics, we next considered how our findings translate to the dynamics on NTRs in a system containing many FG domains. Understanding how “patchy” NTRs and cargoes travel through dense polymeric mediums, such as in the NPC, is an open problem (see [43−45] for related work). Such considerations are faciliated by available experimental data concerning NTF2 diffusing in a bulk solution of F6A [45]. Specifically, we model one NTR diffusing in a bulk solution (polymer melt) of synthetic FG sequences (F1A -F6A) at a physiologically relevant packing fraction (≈0.1), [5, 22]. To verify that the diffusion measurements can be solely attributed to the movements of the NTR particle relative to the polymer melt, and not any co-diffusing polymers that could be stuck to it, we tested whether specific polymers showed a significantly higher number of binding events to the particle (as would be the case if particular polymers were stuck to the particle). We observed that the NTR particle uniformly sampled the polymer binding sites, thus ensuring that particular polymers were not stuck to the particle (see SI Figure 4).

Having demonstrated that we could measure the diffusion of NTRs with respect to a polymer melt, we next investigated the dynamics of the patchy and uniform NTF2 model particles. Somewhat intriguingly, the instantaneous diffusion coefficients for the patchy NTF2 particle are typically about 4-fold larger (Figure 4a) than for the uniform particle, in spite of having − for similar *K*_*D*_s - lower *k*_off_ s in the single-molecule simulations (Fig. 2c). This may be related to the results in Figure 3b, which showed that the presence of a larger number of FSFG motifs (18 for the Nsp1-like sequence instead of 1-6 for the synthetic sequences) resulted in significantly tighter binding for the uniform NTF2 model. Indeed, we observed a two-fold increase in the number of individual - FSFG-particle − binding events for the uniform particle as compared with the particle with 2 patches, which we attribute to the spreading out of NTR affinity through increased numbers of weak binding sites (see SI Figure 4). This implies that, for the polymer densities and polymer-particle size ratios explored here, there is a dynamical cost in binding to large numbers of FSFG motifs in the melt that is analogous to increased friction/drag on the particle, and could provide a physical reason as to why NTRs are not totally covered in binding sites.

**FIG. 4.**
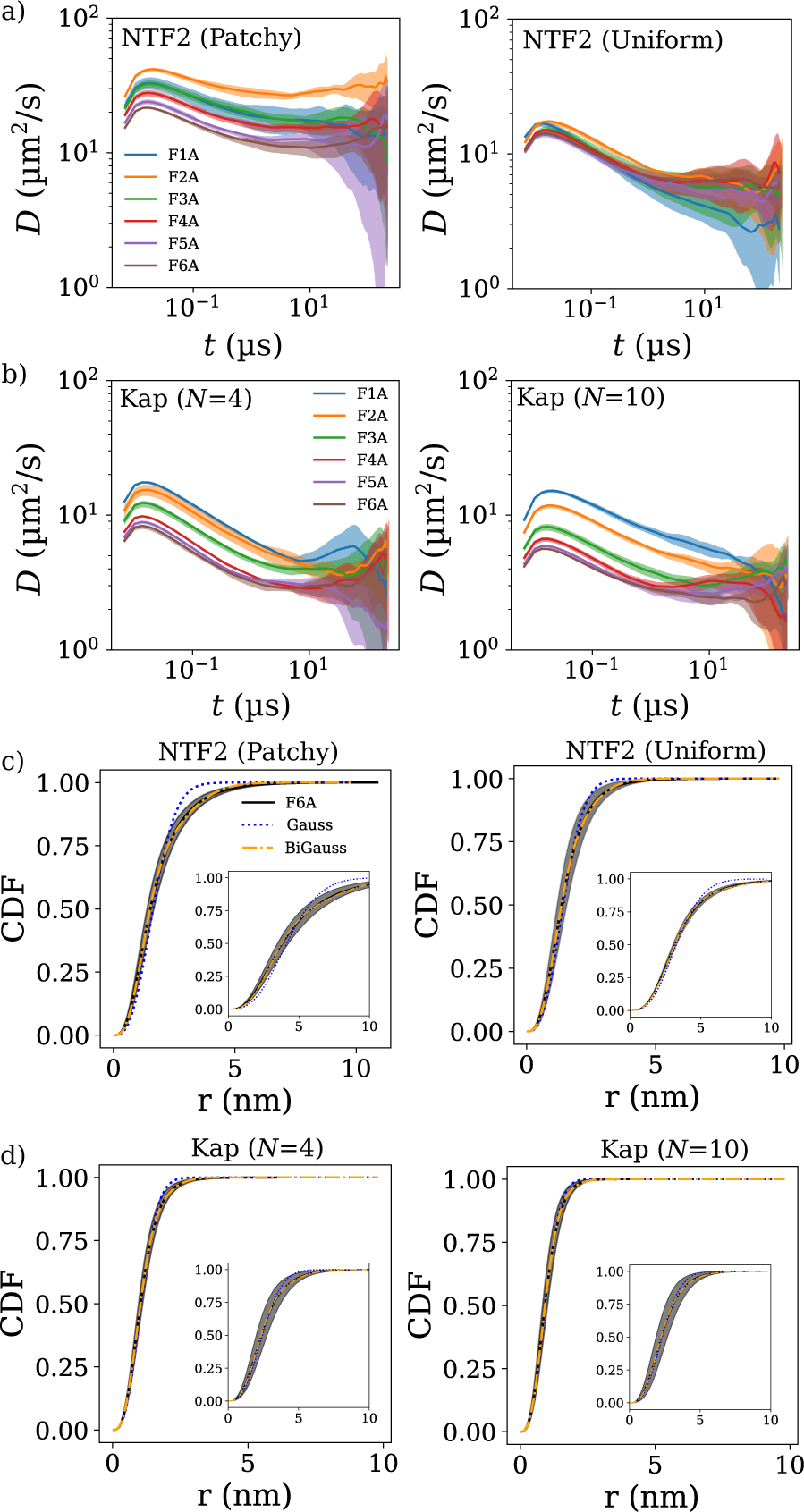
Equilibrium dynamics of a single nuclear transport receptor in a bulk solution of various synthetic FG protein sequences. a) Instantaneous diffusion coefficients for patchy (left) and uniform (right) NTF2 particles, with equivalent single-molecule dissociation constants for the F1A-F6A sequences, as a function of (lag) time. Bands are standard deviations from 5 independent runs. b) Same as (a) but for the Kap model with *N* = 10 and 4 with respective cohesion strengths of **ϵ**′ = 8.0 *k*_*B*_ *T* and 9.0 *k*_*B*_ *T* that give equivalent dissociation constants for the short F1A-F6A sequences. c) Cummulative distribution functions (CDF) for patchy and uniform NTF2 for the selected sequence (F6A) alongside two fits: one using a single mobility model (Gauss) and the other a two mobility model (BiGauss). d) Same as (c) but for the Kap models. For both (c) and (d) the CDFs are for *t ≈* 10^−2^ *s* (main panels) and *t ≈* 10^0^ *s* (inset panels).

As for NTF2, we find for Kap95 that a higher *k*_off_ in the single-molecule simulations (Figure 3c,d) does not translate into substantially faster diffusion in the polymer melt, as illustrated for the Kap95 models for *N* = 4 and *N* = 10 patches with matching single-molecule *K*_*D*_s (Figure 4b). For both Kap particles we observe that their dynamics become slower (≈2 − 3-fold difference) upon increasing the number of cohesive polymer blocks from 1 to 6 and they tend to be slower than patchy NTF2, but are comparable to the uniform NTF2 model. Satisfyingly, the diffusion coefficients for the NTR models (∼10 μm^2^*/*s ≡ 10^−7^ cm^2^*/*s) are in general agreement with previous experimental data (at similar FG densities as in the NPC) [11, 45 − 47]. It appears that with an appropriately chosen single-molecule *K*_*D*_, the unbinding kinetics does not seem to matter too much for the diffusion in the melt. However, the unbinding kinetics might matter more at the peripheries of the NPC, in which faster off-rates could determine faster detachment from the mass of FG Nups occluding the pore (we do not explore this here).

Diffusion in crowded biological environments as well as in crowded polymer melts can often be characterized as anomalous (non-Fickian), arising from the steric, repulsive, and/or attractive interactions between the molecular components [48-50]. Of relevance, many molecules in the cytoplasm and cell nucleus have been shown to diffuse anomalously (subnormally) [51,52]. Since the instantaneous diffusion coefficients decrease with lag time *t* (Figure 4), we infer that the NTRs show subnormal diffusion in the polymer melt consisting of FSFG sequences. This can be further analysed via the cumulative density function CDF(*r, t*), which is the probability of finding a particle - starting at the origin *r* = 0 at time *t* = 0 - at a distance *r* after a time *t*. We next fitted CDF(*r, t*) to a one-mobility and to a two-mobility model, where the latter is defined by two separate ordinary diffusion coefficients [53−57]. The two-mobility provides more degrees of freedom and may therefore expected to better fit the data If that is the case, it should be regarded as an indication of deviations from predictions for normal diffusion, not necessarily as an indication for the presence of two discrete modes of diffusion. Indeed, as expected based on the variation of the instantaneous diffusion coefficients with lag time in Figure 4a, the computed C Fs are better fitted with a two-mobility, iGauss, anomalous diffusion model (with two parameters) than with a simple one-mobility, Gauss model (with one parameter) that applies for normal diffusion (see Figures 4(c,d) and SI for details).

Interestingly, we find that the patchy NTF2 deviates most from a one-mobility model but can be described well using a two-mobility model with a fast diffusion coefficient (*D*_1_) that is ≈ 4 − 5 times greater than the slower one (*D*_2_), with both fast and slow diffusion playing significant roles (35 % fast, 65% slow) (see SI Table 2). Remarkably, the predicted (at *t* = 1 μs) fast (*D*_1_ = {25.7, 21.7} μm^2^/s) and slow (*D*_2_ = {5.5, 5.1} μm^2^/s) diffusion coefficients for patchy and uniform NTF2 particles respectively lie close to the experimentally determined diffusion coefficients (*D*_free_= 23 ± 10, *D*_bound_= 5.6 ± 2.2} μm^2^/s) as found for an NTF2 particle diffusing between a solution of F6A sequences (relating to *D*_bound_), and a reservoir of inert polymers (relating to *D*_free_) [45]. Also, the predominance of the slower diffusion (*>* 60%) observed in our simulations is also corroborated by the predominance of bound diffusion in the same experiments. Specifically, we may interpret the faster diffusion coefficient *D*_1_ to represent unbound diffusion, when the NTF2 particle is free of FSFG motifs, and the slower diffusion coefficient *D*_2_ represents bound diffusion, with the caveat - mentioned above - that this is unlikely to be a sharp distinction.

Expectedly, the two Kap particles (*N* = {4, 10}) did not show significantly different behaviour, as is observed in the instantaneous diffusion coefficients, and showed minor deviations from the one-mobility model. For both Kaps (*N* = {4, 10}) we found that the slow diffusion coefficient *D*_2_ is similar in value to the one-mobility diffusion coefficient, with a large portion (≈ 70 − 80%) of the diffusion arising from *D*_2_, but still a non-negligible portion of faster diffusion (*D*_1_) arose from the two-mobility fits. We note that the size could also be contributing to the slower diffusion (as compared with the NTF2 models). In all cases, we observe that the diffusion coefficients for the mobility models are largely consistent with the independently calculated instantaneous diffusion coefficients. This analysis highlights that the dynamics of NTRs in a heterogenous melt could potentially involve a range of diffusion types, including fast (free/unbound) and slow (bound) diffusion.

## CONCLUSION

In summary, we have presented a physical picture of patterned FG sequences interacting with heterogeneous NTRs. Firstly, our minimal coarse-grained model is in excellent agreement with recent experimental binding data on the level of single-molecules, by only fitting one interaction parameter [13]. We also found that this agreement was maintained even when the surface heterogeneity of the NTR was ignored. This is consistent with our previous observations on NTR uptake in surface-grafted FG domain assemblies as described by mean-field homopolymer models [21,22], and provides a foundation for other computational studies that made similar assumptions [7, 19, 25, 26, 58, 59]. However, we found the uniform NTR model exhibited much faster binding kinetics (with a high “off” rate ∼ 10^6^ s^−1^ and “on” rate ∼ 10^9^ M^−1^s^−1^) than the -patch model for the same dissociation constant, owing to a greater spread of the cohesion over the particle. This suggests that care must be taken when inferring kinetic properties from homopolymer type modelling.

Using analytical theories, one based on a sequence-specific polymer theory and the other based on a statistical mechanical lattice binding model, we further confirmed the roles that sequence heterogeneity of the FG sequences have on FG-NTR binding that the overall number of cohesive blocks largely dictates binding affinity with variations in the exact distances between those blocks being of lesser importance. These theories provide a physical picture and mathematical framework to understand single-molecule binding of patterned polymers and heterogeneous nanoparticles.

Furthermore, we made predictions on the binding of a more realistic FG sequence, based on Nsp1, with NTRs, based on Kap95, having more complex surface patterning. We found that an NTR that is larger than NTF2 but with more cohesive patches resulted in a similar overall binding affinity to the FG sequence, highlighting a possible reason for why larger cargoes, *e*.*g*, importin-*β*, have more FG binding grooves/spots to compensate for its size [10]. Overall, the number of binding patches on the NTR is critical we observe a change from ∼1 mM to ∼10 μ M in the FSFG-NTR binding affinity upon doubling the number of particle cohesive patches.

Our results have potential implications on the large differences between single-molecule *K*_*D*_s as inferred from various experiments, including FSFG constructs and native FG Nups, with values including ∼1 mM [12, 13], ∼10 μ M[11, 14, 15], and ≲ 10 μ [15 − 18]. We reconcile these affinity differences through varying degrees of multivalency, where moderate increases in available binding sites results in large increases in affinity. As such, care must taken in inferring single-molecule binding affinities from macroscopic systems where collective effects, such as the non-linear dependence of dissocation on the binding site concentration, will carry over into any extrapolated single-molecule binding constant.

Lastly, through exploring the diffusion of an NTR in an FG polymer melt we found that the observations from single-molecule kinetics do not necessarily carry over to macroscopic systems. Specifically we did not observe an increase in macroscopic diffusion coefficient upon spreading out the cohesion on an NTRs surface, through more patches, despite this leading to higher *k* s in the single-molecule simulations. For the NTRs explored here, we find generally fast (∼1 − 50 μm^2^*/*s) diffusion in the FSFG polymer melt in accordance with previous experimental data [11, 45 − 47]. Our findings suggest that the macroscopic diffusion of NTRs could be due to a combination of fast and slow modes, with the important caveat that there are many different physical regimes that affect diffusion and it is not known which one is directly relevant to the NPC. Whilst we have not explored the precise microscopic mechanism of NTR motion in an FG polymer melt, we expect that our macroscopic picture can be fully reconciled with microscopic pictures of motion such as the “slide-and-exchange” mechanism [43]. We speculate that whilst the overall diffusion in a dense melt is not greatly affected by changes in the surface properties of the NTR, they may be of particular importance at the nucleoplasmic and cytoplasmic entry points of the NPC where high specificity (*K*_*D*_ and *k*_on_) governs uptake and fast unbinding kinetics (*k*_off_) governs release (although cargo transport may not need the NTRs to be released from the NPC).

Overall, we have presented a minimal yet adaptable physical framework that sets a solid foundation to further explore specific physical questions regarding selective transport in the NPC whilst also elucidating future design principles for the components of artificial transport nanomachines.

## AUTHOR CONTRIBUTIONS

L.K.D. designed the model, developed the theory, performed and analysed the simulations and calculations, intepreted the results, and wrote the manuscript. A. Š. advised on MD simulations, analysis, and interpreted the results. B.W.H. and A.Z. conceived the study, helped design the model and theory, interpreted the results, supervised the project and co-wrote the manuscript. All authors provided feedback on the manuscript.

## ACKNOWLEDGMENTS

We thank Tiantian Zheng, the Hoogenboom group, and other members of the Zilman group for discussions. L.K.D. acknowledges the biophysics research computing cluster at UCL that was used to perform the simulations and analysis. This work was funded by the Royal Society (UF160266 A.Š.) and the UK Engineering and Physical Sciences Research Council (EP/L504889/1, L.K.D. and B.W.H.). A.Z. acknowledges the support of the National Science and Engineering Research Council of Canada (NSERC) through Discovery Grant RGPIN 402591. The authors declare no competing interests.

## Notes

### Competing Interest Statement

The authors have declared no competing interest.

