## Supplementary_Information for "Physical modelling of multivalent interactions in the nuclear pore complex"

(Dated: September 30, 2020)

### CONTENTS

I. Calculating the dissociation constant $K_D$ and off-rate $k_{\text{off}}$ from simulations 2

II. Calculating the instantaneous diffusion coefficient from simulation 3

III. Calculating the cumulative density function from simulations and derivation of the mobility models 3

IV. Analytical calculation for sequence-specific dissociation constants 4
A. Derivation of the second virial coefficient $B_2$ 4
B. Mathematical details for pair-potential factors in $B_2$ 9
C. Mathematical details for incorporating intra-molecular interactions 10

V. Mathematical derivation of $K_D$ for the lattice binding model 14

VI. Supplementary tables and figures 19

References 24

### I. CALCULATING THE DISSOCIATION CONSTANT $K_D$ AND OFF-RATE $k_{\text{off}}$ FROM SIMULATIONS

The dissociation constant (in units of mol·nm$^{-3}$) between one patterned polymer, defined by a sequence of cohesive and non-cohesive blocks, and one patchy-particle was computed using the relation [1]

$$K_D = \frac{N_0}{N_A N_1 (V - V_D)} \tag{1}$$

where $N_A$ is Avogadro's constant, $N_1$ is the number of occurences of at least one cohesive bead on the patterned polymer being in the cohesive range $2r_0$ of a cohesive bead on a patchy-particle (defining the bound state), $N_0$ is the number of occurences where no cohesive beads on the patterned polymer are in the cohesive range $2r_0$ of a cohesive bead on a patchy-particle (defining the unbound state), and $V_D$ is the dimerization volume approximated as

*

$\frac{4}{3}\pi(N+N_m)(2r_0)^3$, where $N$ and $N_m$ are the number of particle patches and cohesive polymer beads respectively. To compare with experimental data the calculated $K_D$ is converted to molar units (M) using 1 nm$^{-3} \equiv 10^{-24}$ liters. Note that for simplicity we use the sum of all spherical volumes, and do not correct for any overlap in the potential ranges between beads. To calculate the dissociation constant between a patterned polymer and uniform particle, the volume of the spacer bead comprising the uniform particle is used (see supplementary table 1 for the relevant $r_0$).

To calculate the off-rate we use the standard definition $k_{\text{off}} = t_{\text{res}}^{-1}$, where $t_{\text{res}}$ is the time spent in the bound state. The definition of the bound state is the same as is used for the $K_D$ calculation. In practice the residence time is computed as $t_{\text{res}} = n_{\text{bound}}\delta t$ where $n_{\text{bound}}$ is the number of consecutive MD frames that the two molecules are in the bound state and $\delta t$ is the simulation timestep.

### II. CALCULATING THE INSTANTANEOUS DIFFUSION COEFFICIENT FROM SIMULATION

First the time-averaged mean squared displacement $MSD$ was computed using the freud package [2] and calculated according to $MSD(m) = \frac{1}{N_t-m}\sum_{k=0}^{N_t-m-1}(\mathbf{r}(k+m)-\mathbf{r}(k))^2$, where $m$ denotes the current discrete simulation time, $N_t$ is the total number of time-frames, $k$ is an index over the time-frames, and $\mathbf{r}$ is the position of the particle. The instantaneous diffusion coefficient is then given as $D(t) = MSD(t)/6t$ where the time is calculated as $t = m\delta t$.

### III. CALCULATING THE CUMULATIVE DENSITY FUNCTION FROM SIMULATIONS AND DERIVATION OF THE MOBILITY MODELS

The cumulative density function $CDF(r,t)$ is the probability that a particle, starting from the origin, is within a sphere of radius $r$ at time $t$. We calculated $CDF(r,t)$ by counting the number of absolute displacements $|\mathbf{r}(t_{i+n}) - \mathbf{r}(t_i)| < r$, where $n$ is the discrete time interval, in a single trajectory and normalizing the count by the total number of considered time points, we then averaged the result over five independent runs (based on a similar procedure [3]). To characterize the type of diffusion as observed in simulations, we fit the calculated CDFs to two mobility models. One model is based on normal diffusion defined (in $d$ dimensions) by a probability distribution $P(r,t)$ given as

$$P(r,t) = (4\pi Dt)^{-d/2} \exp\left(-\frac{r^2}{4Dt}\right), \tag{2}$$

where $D$ is the diffusion coefficient. The $CDF(r,t)$ is then

$$CDF(r,t) = \int_0^r P(r,t)dV_d(r), \tag{3}$$

where the 'volume' element is $dV_d(r) = \frac{2\pi^{d/2}r^{d-1}}{\Gamma(d/2)}dr$ and $\Gamma()$ is the Gamma function. Setting $d = 3$ and integrating results in

$$CDF(r,t) = \text{erf}\left(\frac{r}{2\sqrt{Dt}}\right) - \frac{r}{\sqrt{\pi Dt}} \exp\left(-\frac{r^2}{4Dt}\right), \tag{4}$$

where erf($x$) is the error function. The two mobility model is defined by two diffusion coefficients $D_1$ and $D_2$, where $D_1 > D_2$, of defined weighting $0 \leq w \leq 1$ such that the resulting model is

$$\begin{aligned} CDF(r,t) = (1-w)&\left[\text{erf}\left(\frac{r}{2\sqrt{D_1 t}}\right) - \frac{r}{\sqrt{\pi D_1 t}} \exp\left(-\frac{r^2}{4D_1 t}\right)\right] \\ w&\left[\text{erf}\left(\frac{r}{2\sqrt{D_2 t}}\right) - \frac{r}{\sqrt{\pi D_2 t}} \exp\left(-\frac{r^2}{4D_2 t}\right)\right]. \end{aligned} \tag{5}$$

### IV. ANALYTICAL CALCULATION FOR SEQUENCE-SPECIFIC DISSOCIATION CONSTANTS

#### A. Derivation of the second virial coefficient $B_2$

To further elucidate the binding affinity between a patterned polymer ($A$) and a particle ($B$), we used an analytical theory, based upon a cluster expansion-variational method, that takes into account sequence heterogeneity [4, 5].

Motivated by [4], the theory is derived in its most general form where the patterned polymer and particle consist of, respectively, $N^A$ and $N^B$ constituent monomers of which the coordinates are defined through $\mathbf{R}^A = \{\mathbf{r}_1^A, \ldots, \mathbf{r}_{N^A}^A\}$ and $\mathbf{R}^B = \{\mathbf{r}_1^B, \ldots, \mathbf{r}_{N^B}^B\}$, respectively. Later on, $N^B = 1$ will be imposed to simplify the calculation and to replicate the uniform particle model as described in the main text. The patterning of the polymer and particle are defined, respectively, through $S^A = \left\{s_1^A, \ldots, s_{N^A}^A\right\}$ and $S^B = \left\{s_1^B, \ldots, s_{N^B}^B\right\}$, where $s = 0, 1$ with 0 and 1 representing non-cohesive and cohesive beads. When $N^B = 1$ we set $S^B = \{1\}$.

The binding affinity between the two molecules $A$ and $B$ is characterized through the dissociation constant $K_D$ (in units of mol·nm$^{-3}$) that can be defined – in the dilute limit – as

$$K_D = \frac{(1 - P_{AB})}{P_{AB}} \frac{1}{N_A V}, \tag{6}$$

where $P_{AB} \equiv P_{BA}$ is the binding probability, $V$ is the system volume, and $N_A$ is Avogradro's number [1, 4, 6]. The unbound probability, $1 - P_{AB}$, can be written as

$$1 - P_{AB} = \frac{V \mathcal{Q}_A \mathcal{Q}_B}{\mathcal{Q}_{AB}}, \tag{7}$$

where $\mathcal{Q}_i$, $i = A, B, AB$, are the configurational partition functions [4] defined as

$$\begin{aligned}
\mathcal{Q}_i &= \frac{1}{V} \int \mathcal{D}[\mathbf{R^i}] e^{-\beta \mathcal{H}^i[\mathbf{R}^i]}, \quad i = A, B, \\
\mathcal{Q}_{AB} &= \frac{1}{V} \int \mathcal{D}[\mathbf{R^A}] \mathcal{D}[\mathbf{R^B}] e^{-\beta(\mathcal{H}^A[\mathbf{R}^A] - \mathcal{H}^B[\mathbf{R}^B] - \mathcal{U}^{AB}[\mathbf{R}^A, \mathbf{R}^B])},
\end{aligned} \tag{8}$$

where $\int \mathcal{D}[\mathbf{R}^i] \equiv \int \prod_{s=1}^{N^i} d\mathbf{r}_s^i$, with $i = A, B$, is an integration over all configurations, $\beta = 1/k_B T$, $\mathcal{H}^i[\mathbf{R}^i]$ $(i = A, B)$ is the Hamiltonian, and $\mathcal{U}^{AB}[\mathbf{R}^A, \mathbf{R}^B]$ is the intermolecular interaction energy between $A$ and $B$ given as

$$\mathcal{U}^{AB}[\mathbf{R}^A, \mathbf{R}^B] = \sum_{a=1}^{N^A} \sum_{b=1}^{N^B} \left(\mathcal{V}_{ab}^{AB}\left(r_{ab}^{AB}\right)\right), \tag{9}$$

where $r_{ab}^{AB} = |\mathbf{r}_a^A - \mathbf{r}_b^B|$ is the distance between monomer $a$ on $A$ and monomer $b$ on $B$ and $\mathcal{V}_{ab}^{AB}$ is the interaction energy between those monomers [4]. Hence, making use of the relation for the second order virial coefficient $B_2 = V - \mathcal{Q}_{AB}/\mathcal{Q}_A \mathcal{Q}_B$, one obtains [4, 7]

$$K_D = \frac{V \mathcal{Q}_A \mathcal{Q}_B}{\mathcal{Q}_{AB}} \left(1 - \frac{V \mathcal{Q}_A \mathcal{Q}_B}{\mathcal{Q}_{AB}}\right)^{-1} \frac{1}{N_A V} = \frac{V/(V - B_2)}{1 - (V/(V - B_2))} \frac{1}{N_A V} = -\frac{1}{N_A B_2}. \tag{10}$$

Using equation 8, the second virial coefficient can also be written as

$$B_2 = V \int \mathcal{D}[\mathbf{R}^A] \mathcal{D}[\mathbf{R}^B] \mathcal{P}^A[\mathbf{R}^A] \mathcal{P}^B[\mathbf{R}^B] \left(1 - \exp(-\beta \mathcal{U}^{AB}[\mathbf{R}^A, \mathbf{R}^B])\right), \tag{11}$$

where $\mathcal{P}^i[\mathbf{R}^i]$ is the probability distribution for molecule $i$. The cluster expansion of $\exp(-\beta\mathcal{U}^{AB}[\mathbf{R}^A,\mathbf{R}^B])$ is given by

$$\begin{aligned}
\exp(-\beta\mathcal{U}^{AB}[\mathbf{R}^A,\mathbf{R}^B]) &= \exp(-\beta\sum_{a=1}^{N^A}\sum_{b=1}^{N^B}\mathcal{V}_{ab}^{AB}\left(r_{ab}^{AB}\right)),\\
&= \prod_{a=1}^{N^A}\prod_{b=1}^{N^B}\left(f_{ab}+1\right),\\
&= 1+\sum_{a=1}^{N^A}\sum_{b=1}^{N^B}f_{ab}+\sum_{a'\geqslant a=1}^{N^A}\sum_{b'\geqslant b=1}^{N^B}f_{a'b'}f_{ab}-\sum_{a=1}^{N^A}\sum_{b=1}^{N^B}f_{ab}^2,\\
&+\mathcal{O}\left(f^3\right)
\end{aligned}\tag{12}$$

where $f_{ab}\equiv\exp(-\beta\mathcal{V}_{ab}^{AB}\left(r_{ab}^{AB}\right))-1$ is the Mayer function [4]. One can introduce the Fourier representation of the Mayer function as

$$f_{ab}=\int\frac{d^3k}{(2\pi)^3}\left[\hat{f}(\mathbf{k})\right]_{ab}\exp(i\mathbf{k}\cdot\mathbf{r}),\tag{13}$$

where $\left[\hat{f}(\mathbf{k})\right]_{ab}$ is an element of an $N^A\times N^B$ matrix. Substituting equation 12 into equation 11 results in

$$B_2=-\sum_{a=1}^{N^A}\sum_{b=1}^{N^B}\left[\hat{f}(\mathbf{0})\right]-\frac{1}{2}\int\frac{d^3k}{(2\pi)^3}\mathrm{Tr}\left[\hat{f}(\mathbf{k})\hat{P}^B(-\mathbf{k})\hat{f}(\mathbf{k})^T\hat{P}^A(-\mathbf{k})-\hat{f}(\mathbf{k})\hat{f}^T(-\mathbf{k})\right],\tag{14}$$

where Tr $[\ldots]$ denotes the trace (double sum over indices), "T" superscript is the transpose, and $\left[\hat{P}^i(\mathbf{k})\right]_{ab}=\int\mathcal{D}[\mathbf{R}^i]\mathcal{P}^i[\mathbf{R}^i]\exp(i\mathbf{k}\cdot(\mathbf{r}_a^i-\mathbf{r}_b^i))$ is the Fourier transform of the intra-molecular monomer-monomer correlation function. The detailed derivation from equation 12 to equation 14 is explicitly given in the supplementary information of [4].

The attractive interaction between cohesive bead $a$ on $A$ and $b$ on $B$ is implemented by the Morse potential

$$\beta\mathcal{V}_{ab}^{ij}(r_{ab}^{ij})=\epsilon^{ij}s_a^is_b^j\left(\exp\left(-2\alpha(r_{ab}^{ij})\right)-2\exp\left(-\alpha(r_{ab}^{ij})\right)\right),\tag{15}$$

where $\epsilon^{ij}$ is the dimensionless cohesive energy between a cohesive monomer on molecule $i$ and a cohesive monomer on molecule $j$ and $\alpha$ is the decay length. As a starting point, the molecules $A$ and $B$ are first treated as ideal-chains of Kuhn length $l_i$ leading to correlation functions

$$\left[\hat{P}^i(\mathbf{k})\right]_{a'a} \approx \exp\left(-\frac{(kl_i)^2|a'-a|}{6}\right), \tag{16}$$

where $i = A, B$. We note that when we later set $N^B = 1$ we will use $\left[\hat{P}^B(\mathbf{k})\right]_{a'a} = 1$. After performing a high temperature expansion of the Mayer-function (see below), the second virial coefficient, $B_2$, is then given by

$$\begin{aligned} B_2 = -\frac{15\pi\epsilon^{AB}}{\alpha^3}\sum_{a=1}^{N^A}\sum_{b=1}^{N^B} s_a^A s_b^B - \\ \int \frac{576\alpha^6(\epsilon^{AB})^2k^2\left(5\alpha^2+2k^2\right)^2}{\left(4\alpha^4+5\alpha^2k^2+k^4\right)^4} \sum_{a',a=1}^{N^A}\sum_{b',b=1}^{N^B} s_{a'}^A s_a^A s_{b'}^B s_b^B \\ \exp\left(-\frac{(kl_A)^2|a'-a|}{6}\right)\exp\left(-\frac{(kl_B)^2|b'-b|}{6}\right)dk, \end{aligned} \tag{17}$$

with the first term being the cohesive mean field contribution and the second term accounting for sequence-specific intermolecular cohesive correlations. However, equation 17 does not yet account for intra-molecular excluded volume and cohesive interactions, in order to do so a variational approach was used, as previously done in [5].

First the Hamiltonian for a polymer whose $N$ monomers interact with each other through excluded volume and cohesion is defined as

$$\begin{aligned} \beta H_T = \frac{3}{2l}\int_0^L d\tau\left(\frac{d\mathbf{r}(\tau)}{d\tau}\right)^2 + l\int_0^L d\tau\int_0^\tau d\tau'\omega(\tau,\tau')\delta\left[r(\tau)-r(\tau')\right] + \\ \frac{\epsilon}{l^2}\int_0^L d\tau\int_0^\tau d\tau' s_\tau s_{\tau'}\left(\exp\left(-2\alpha(|\mathbf{r}_\tau-\mathbf{r}_{\tau'}|)\right) - 2\exp\left(-\alpha(|\mathbf{r}_\tau-\mathbf{r}_{\tau'}|)\right)\right), \end{aligned} \tag{18}$$

where $\tau$ is the contour variable along the polymer, $L = Nl$ is the contour length, and $\omega(\tau,\tau')$ is the dimensionless excluded volume parameter. The Hamiltonian $H_T$, describing all the intra-molecular interactions, can be mapped to an ideal-chain Hamiltonian $H'$ with a renormalized Kuhn length $l'$ written as

$$\beta H' = \frac{3}{2l'}\int_0^L d\tau\left(\frac{d\mathbf{r}(\tau)}{d\tau}\right)^2, \tag{19}$$

where one imposes that the mapping does not change the configurational properties, *e.g.*, the radius of gyration $R_G$, up to first perturbative order in $H' - H_T$. After performing this re-normalization procedure, of which the details can be found below, one obtains an equation for the ratio of the renormalized Kuhn length for monomers $m$ and $n$ on the same polymer to the actual Kuhn length defined as $x_{m,n} = l'_{m,n}/l$.

In order to determine $x_{m,n}$ through equation 36 one must first determine the monomer-monomer dimensionless excluded volume parameter $\omega_{m,n}$ (defined in equation 20 below). $\omega_{m,n}$ is first determined by assuming that it does not depend on the sequence (homo-polymer) and then by switching off intramolecular cohesion, i.e., $\epsilon = 0$, resulting in

$$\omega_{m,n} = \frac{2\sqrt{\frac{2}{3}}\pi^{3/2}\,(n-m)\,(x_{m,n}-1)}{3lx_{m,n}\Omega\sqrt{\frac{1}{(lx_{m,n})^5}}}, \tag{20}$$

where $\Omega$ is defined as

$$\Omega = \sum_{q=m+1}^{M}\left(\sum_{p=n}^{m}\frac{(m-p)^2}{(q-p)^{5/2}}\right) + \sum_{q=m+1}^{M}\left(\sum_{p=1}^{n-1}\frac{(m-n)^2}{(q-p)^{5/2}}\right) + \sum_{q=n}^{m}\left(\sum_{p=1}^{n-1}\frac{(q-n)^2}{(q-p)^{5/2}}\right) + \sum_{q=n+1}^{m}\left(\sum_{p=n}^{q-1}\frac{1}{\sqrt{q-p}}\right). \tag{21}$$

Note that this only applies to molecule A, *i.e.*, the patterned polymer, and not molecule B (the particle) as it will be assumed B consists of one bead only. The parameterization procedure for molecule A is as follows: (i) First simulate a polymer with no intra-molecular cohesion (corresponding to sequence F1A) and calculate its radius of gyration. (ii) Compute the radii of gyration for various values of the averaged excluded volume parameter $\langle\omega\rangle$ using $R_G^2 = 1/N^2 \sum_{m>n}\langle(\mathbf{r}_m - \mathbf{r}_n)^2\rangle = |m-n|l^2x_{m,n}$, finding the $\langle\omega\rangle$ that reproduces the $R_G$ from simulation. (iii) Then simulate polymers with different number of cohesive blocks (sequences F1A - F6A) and calculate their radii of gyration. (iv) Calculate $R_G$'s from theory using $\omega_{m,n} = \langle\omega\rangle$ in equation 36 in addition to switching on cohesion, i.e., $|\epsilon| > 0$, to numerically determine the best match intra-molecular cohesion parameter $\epsilon'$. (v) Finally, determine each $x_{m,n}$. The resulting second virial coefficient, after setting $N^B = 1$, then becomes

$$B_2 = -\frac{15\pi\epsilon^{AB}}{\alpha^3}\sum_{a=1}^{N^A} s_a^A - \int \frac{576\alpha^6(\epsilon^{AB})^2k^2\left(5\alpha^2+2k^2\right)^2}{\left(4\alpha^4+5\alpha^2k^2+k^4\right)^4}\sum_{a',a=1}^{N^A} s_{a'}^A s_a^A \exp\left(-\frac{x_{a',a}^A(kl_A)^2|a'-a|}{6}\right)dk, \quad (22)$$

where the first term is the mean field contribution and the second term is the contribution from monomer-monomer correlations. Equations 22 and 10 are then used to determine the $K_D$'s. The subsections below provide explicit mathematical details regarding the cohesive factors in $B_2$ and in determining $x_{a',a}^A$ above.

### B. Mathematical details for pair-potential factors in $B_2$

Fourier transforming equation 15 and integrating using spherical coordinates gives

$$\begin{aligned}\left[\hat{\mathcal{V}}(\mathbf{k})\right]_{ab} =& \epsilon^{ij}s_a^i s_b^j \int_{r_0}^{\infty}\int_0^{\pi}\int_0^{2\pi}\left(\exp\left(-2\alpha(r_{ab}^{ij}-r_0^{ij})\right)-2\exp\left(-\alpha(r_{ab}^{ij}-r_0^{ij})\right)\right), \\ & \exp\left(i\mathbf{k}\cdot r_{ab}^{ij}\cos(\phi)\right)\sin(\phi)(r_{ab}^{ij})^2 d\theta d\phi dr_{ab}^{ij} \\ =& -\frac{48\pi\epsilon^{ij}s_a^i s_b^j\left(5\alpha^5+2\alpha^3k^2\right)}{\left(4\alpha^4+5\alpha^2k^2+k^4\right)^2}, \qquad (23)\end{aligned}$$

with $k \equiv |\mathbf{k}|$. Next, under the assumption of weak interactions, a high temperature expansion to the Mayer function is performed leading to

$$\begin{aligned}\left[\hat{f}(\mathbf{k})\right]_{ab} =& -\left[\hat{\mathcal{V}}(\mathbf{k})\right]_{ab} + \frac{1}{2}\left[\hat{\mathcal{V}}(\mathbf{k})\right]_{ab}^2 + \ldots \\ =& (\epsilon^{ij}s_a^i s_b^j)\left(\frac{240\pi\alpha^5}{\left(4\alpha^4+5\alpha^2k^2+k^4\right)^2} + \frac{96\pi\alpha^3k^2}{\left(4\alpha^4+5\alpha^2k^2+k^4\right)^2}\right) + \\ & +(\epsilon^{ij}s_a^i s_b^j)^2\left(\frac{28800\pi^2\alpha^{10}}{\left(4\alpha^4+5\alpha^2k^2+k^4\right)^4} + \frac{23040\pi^2\alpha^8k^2}{\left(4\alpha^4+5\alpha^2k^2+k^4\right)^4} + \frac{4608\pi^2\alpha^6k^4}{\left(4\alpha^4+5\alpha^2k^2+k^4\right)^4}\right) \\ & + \ldots \qquad (24)\end{aligned}$$

Referring to equation 14, keeping terms of $-\frac{1}{2}\frac{1}{(2\pi)^3}\left[\hat{f}(\mathbf{k})\right]_{ab}^2$ up to $\mathcal{O}((\epsilon^{ij}s_a^i s_b^j)^2)$ gives

$$-\frac{1}{2}\frac{1}{(2\pi)^3}\left[\hat{f}(\mathbf{k})\right]^2_{ab} = -\frac{144\alpha^6(\epsilon^{ij}s^i_a s^j_b)^2\left(5\alpha^2+2k^2\right)^2}{\pi\left(4\alpha^4+5\alpha^2k^2+k^4\right)^4} + \mathcal{O}((\epsilon^{ij}s^i_a s^j_b)^3) + \ldots, \tag{25}$$

and integrating using spherical coordinates leads to

$$\begin{aligned}\int_0^\pi\int_0^{2\pi} -\frac{144\alpha^6(\epsilon^{ij}s^i_a s^j_b)^2k^2\left(5\alpha^2+2k^2\right)^2\sin(\phi)}{\pi\left(4\alpha^4+5\alpha^2k^2+k^4\right)^4}d\theta d\phi \\ = -\frac{576\alpha^6(\epsilon^{ij}s^i_a s^j_b)^2k^2\left(5\alpha^2+2k^2\right)^2}{\left(4\alpha^4+5\alpha^2k^2+k^4\right)^4},\end{aligned} \tag{26}$$

which directly enters equation 17.

### C. Mathematical details for incorporating intra-molecular interactions

Let $\chi$ be an observable that stores information about the configuration of the polymer, and whose ensemble average with respect to $H_T$ is given by

$$\langle\chi\rangle_T = \langle\chi\rangle' + \langle\chi\rangle'\langle H' - H_T\rangle' - \langle\chi(H' - H_T)\rangle + \mathcal{O}((H' - H_T)^2), \tag{27}$$

where $\langle\ldots\rangle$ is the ensemble average computed over configuration space. The condition that $\langle\chi\rangle_T = \langle\chi\rangle'$ to first order implies that $\langle\chi\rangle'\langle H'-H_T\rangle' = \langle\chi(H'-H_T)\rangle'$. Here, the square of monomer-monomer distance is used as the variable $\chi = (\mathbf{r}(\tau_2) - \mathbf{r}(\tau_1))^2$ such that

$$\langle(\mathbf{r}(\tau_2) - \mathbf{r}(\tau_1))^2\rangle'\langle H' - H_T\rangle' = \langle(\mathbf{r}(\tau_2) - \mathbf{r}(\tau_1))^2\,(H' - H_T)\rangle', \tag{28}$$

which determines a renormalised Kuhn length for each monomer-monomer pair $l'(\tau_2, \tau_1)$. Placing equations 18 and 19 into equation 28 gives

$$
\begin{aligned}
&\left\langle(\mathbf{r}(\tau_2)-\mathbf{r}(\tau_1))^2\frac{3}{2}(\frac{1}{l'}-\frac{1}{l})\int_0^L d\tau\left(\frac{d\mathbf{r}(\tau)}{d\tau}\right)\right\rangle' - \\
&\qquad\left\langle(r(\tau')-r(\tau))^2\right\rangle'\left\langle\frac{3}{2}(\frac{1}{l'}-\frac{1}{l})\int_0^L d\tau\left(\frac{d\mathbf{r}(\tau)}{d\tau}\right)\right\rangle' \\
&= \left\langle(\mathbf{r}(\tau_2)-\mathbf{r}(\tau_1))^2 l\int_0^L d\tau\int_0^\tau d\tau'\omega(\tau,\tau')\delta\left[\mathbf{r}(\tau)-\mathbf{r}(\tau')\right]\right\rangle' - \\
&\qquad\left\langle(\mathbf{r}(\tau_2)-\mathbf{r}(\tau_1))^2\right\rangle'\left\langle l\int_0^L d\tau\int_0^\tau d\tau'\omega(\tau,\tau')\delta\left[\mathbf{r}(\tau)-\mathbf{r}(\tau')\right]\right\rangle' \\
&+\left\langle(\mathbf{r}(\tau_2)-\mathbf{r}(\tau_1))^2\frac{\epsilon}{l^2}\int_0^L d\tau\int_0^\tau d\tau' s_\tau s_{\tau'}\left(\exp\left(-2\alpha(|\mathbf{r}_\tau-\mathbf{r}_{\tau'}|)\right)-2\exp\left(-\alpha(|\mathbf{r}_\tau-\mathbf{r}_{\tau'}|)\right)\right)\right\rangle' \\
&-\left\langle(\mathbf{r}(\tau_2)-\mathbf{r}(\tau_1))^2\right\rangle'\left\langle\frac{\epsilon}{l^2}\int_0^L d\tau\int_0^\tau d\tau' s_\tau s_{\tau'}\left(\exp\left(-2\alpha(|\mathbf{r}_\tau-\mathbf{r}_{\tau'}|)\right)-2\exp\left(-\alpha(|\mathbf{r}_\tau-\mathbf{r}_{\tau'}|)\right)\right)\right\rangle'.
\end{aligned}
\tag{29}
$$

Using properties of the ideal-chain [5] the left-hand side of equation 29 reduces to

$$
\begin{aligned}
&\left\langle(\mathbf{r}(\tau_2)-\mathbf{r}(\tau_1))^2\frac{3}{2}(\frac{1}{l'}-\frac{1}{l})\int_0^L d\tau\left(\frac{d\mathbf{r}(\tau)}{d\tau}\right)\right\rangle' - \\
&\qquad\left\langle(\mathbf{r}(\tau_2)-\mathbf{r}(\tau_1))^2\right\rangle'\left\langle\frac{3}{2}(\frac{1}{l'}-\frac{1}{l})\int_0^L d\tau\left(\frac{d\mathbf{r}(\tau)}{d\tau}\right)\right\rangle' = l'^2(\tau_2-\tau_1)\left(\frac{1}{l'}-\frac{1}{l}\right),
\end{aligned}
\tag{30}
$$

additionally the excluded volume terms, the first two terms on the right hand side of equation 29, can be written as [5]

$$
\begin{aligned}
&\left\langle(\mathbf{r}(\tau_2)-\mathbf{r}(\tau_1))^2 l\int_0^L d\tau\int_0^\tau d\tau'\omega(\tau,\tau')\delta\left[\mathbf{r}(\tau)-\mathbf{r}(\tau')\right]\right\rangle' - \\
&\qquad\left\langle(\mathbf{r}(\tau_2)-\mathbf{r}(\tau_1))^2\right\rangle'\left\langle l\int_0^L d\tau\int_0^\tau d\tau'\omega(\tau,\tau')\delta\left[\mathbf{r}(\tau)-\mathbf{r}(\tau')\right]\right\rangle' \\
&= -\frac{l'^2}{3}\left(\left(\frac{1}{2\pi}\right)^3\left(\frac{3}{l'}\right)^5\right)^{1/2}\left(\int_{\tau_1}^{\tau_2} d\tau\int_0^{\tau_1} d\tau'\omega(\tau,\tau')\frac{(\tau-\tau_1)^2}{(\tau-\tau')^{5/2}}\right. \\
&+\int_{\tau_1}^{\tau_2} d\tau\int_{\tau_1}^{\tau} d\tau'\omega(\tau,\tau')\frac{1}{(\tau-\tau')^{1/2}}+\int_{\tau_2}^{L} d\tau\int_0^{\tau_1} d\tau'\omega(\tau,\tau')\frac{(\tau_2-\tau_1)^2}{(\tau-\tau')^{5/2}} \\
&\left.+\int_{\tau_2}^{L} d\tau\int_{\tau_1}^{\tau_2} d\tau'\omega(\tau,\tau')\frac{(\tau_2-\tau')^2}{(\tau-\tau')^{5/2}}\right)
\end{aligned}
\tag{31}
$$

The cohesive terms, the last two terms on the right hand side of equation 29, can be simplified also. One can define

$$\exp\left(-2\alpha\mathbf{r}\right) - 2\exp\left(-\alpha\mathbf{r}\right) = 48\pi \int \frac{d^3k}{(2\pi)^3} \frac{\exp(i\mathbf{k}\cdot\mathbf{r})\left(5\alpha^5 + 2\alpha^3k^2\right)}{\left(4\alpha^4 + 5\alpha^2k^2 + k^4\right)^2}, \tag{32}$$

which results in the last two terms of equation 29 being expressed as

$$\begin{aligned}
&\left\langle (\mathbf{r}(\tau_2) - \mathbf{r}(\tau_1))^2 \frac{\epsilon}{l^2} \int_0^L d\tau \int_0^\tau d\tau' s_\tau s_{\tau'} \left(\exp\left(-2\alpha(|\mathbf{r}_\tau - \mathbf{r}_{\tau'}|)\right) - 2\exp\left(-\alpha(|\mathbf{r}_\tau - \mathbf{r}_{\tau'}|)\right)\right) \right\rangle' \\
&-\left\langle (\mathbf{r}(\tau_2) - \mathbf{r}(\tau_1))^2 \right\rangle' \left\langle \frac{\epsilon}{l^2} \int_0^L d\tau \int_0^\tau d\tau' s_\tau s_{\tau'} \left(\exp\left(-2\alpha(|\mathbf{r}_\tau - \mathbf{r}_{\tau'}|)\right) - 2\exp\left(-\alpha(|\mathbf{r}_\tau - \mathbf{r}_{\tau'}|)\right)\right) \right\rangle' \\
&= \left\langle (\mathbf{r}(\tau_2) - \mathbf{r}(\tau_1))^2 \frac{48\pi\epsilon}{l^2} \int_0^L d\tau \int_0^\tau d\tau' s_\tau s_{\tau'} \left( \int \frac{d^3k}{(2\pi)^3} \frac{\exp(i\mathbf{k}\cdot|\mathbf{r}_\tau - \mathbf{r}_{\tau'}|)\left(5\alpha^5 + 2\alpha^3k^2\right)}{\left(4\alpha^4 + 5\alpha^2k^2 + k^4\right)^2} \right) \right\rangle' \\
&-\left\langle (\mathbf{r}(\tau_2) - \mathbf{r}(\tau_1))^2 \right\rangle' \left\langle \frac{48\pi\epsilon}{l^2} \int_0^L d\tau \int_0^\tau d\tau' s_\tau s_{\tau'} \left( \int \frac{d^3k}{(2\pi)^3} \frac{\exp(i\mathbf{k}\cdot|\mathbf{r}_\tau - \mathbf{r}_{\tau'}|)\left(5\alpha^5 + 2\alpha^3k^2\right)}{\left(4\alpha^4 + 5\alpha^2k^2 + k^4\right)^2} \right) \right\rangle'.
\end{aligned} \tag{33}$$

Performing the configurational integrals on the left hand side of equation 33 using an ideal-chain Hamiltonian [5] results in

$$\begin{aligned}
&\left\langle (\mathbf{r}(\tau_2) - \mathbf{r}(\tau_1))^2 \frac{48\pi\epsilon}{l^2} \int_0^L d\tau \int_0^\tau d\tau' s_\tau s_{\tau'} \left( \int \frac{d^3k}{(2\pi)^3} \frac{\exp(i\mathbf{k}\cdot|\mathbf{r}_\tau - \mathbf{r}_{\tau'}|)\left(5\alpha^5 + 2\alpha^3k^2\right)}{\left(4\alpha^4 + 5\alpha^2k^2 + k^4\right)^2} \right) \right\rangle' \\
&-\left\langle (\mathbf{r}(\tau_2) - \mathbf{r}(\tau_1))^2 \right\rangle' \left\langle \frac{48\pi\epsilon}{l^2} \int_0^L d\tau \int_0^\tau d\tau' s_\tau s_{\tau'} \left( \int \frac{d^3k}{(2\pi)^3} \frac{\exp(i\mathbf{k}\cdot|\mathbf{r}_\tau - \mathbf{r}_{\tau'}|)\left(5\alpha^5 + 2\alpha^3k^2\right)}{\left(4\alpha^4 + 5\alpha^2k^2 + k^4\right)^2} \right) \right\rangle' \\
&= -\frac{16\pi\epsilon l'^2}{3l^2} \Bigg( \int_{\tau_1}^{\tau_2} d\tau \int_0^{\tau_1} d\tau' s_\tau s_{\tau'} (\tau - \tau_1)^2 \int \frac{d^3k k^2}{(2\pi)^3} \frac{\left(5\alpha^5 + 2\alpha^3k^2\right)}{\left(4\alpha^4 + 5\alpha^2k^2 + k^4\right)^2} \exp\left(-\frac{k^2 l' |\tau - \tau'|}{6}\right) \\
&\quad + \int_{\tau_1}^{\tau_2} d\tau \int_{\tau_1}^{\tau} d\tau' s_\tau s_{\tau'} (\tau - \tau')^2 \int \frac{d^3k k^2}{(2\pi)^3} \frac{\left(5\alpha^5 + 2\alpha^3k^2\right)}{\left(4\alpha^4 + 5\alpha^2k^2 + k^4\right)^2} \exp\left(-\frac{k^2 l' |\tau - \tau'|}{6}\right) \\
&\quad + \int_{\tau_2}^{L} d\tau \int_0^{\tau_1} d\tau' s_\tau s_{\tau'} (\tau_2 - \tau_1)^2 \int \frac{d^3k k^2}{(2\pi)^3} \frac{\left(5\alpha^5 + 2\alpha^3k^2\right)}{\left(4\alpha^4 + 5\alpha^2k^2 + k^4\right)^2} \exp\left(-\frac{k^2 l' |\tau - \tau'|}{6}\right) \\
&\quad + \int_{\tau_2}^{L} d\tau \int_{\tau_1}^{\tau_2} d\tau' s_\tau s_{\tau'} (\tau_2 - \tau')^2 \int \frac{d^3k k^2}{(2\pi)^3} \frac{\left(5\alpha^5 + 2\alpha^3k^2\right)}{\left(4\alpha^4 + 5\alpha^2k^2 + k^4\right)^2} \exp\left(-\frac{k^2 l' |\tau - \tau'|}{6}\right) \Bigg).
\end{aligned} \tag{34}$$

Then making use of the identity

$$\int \frac{d^3k k^2}{(2\pi)^3} \frac{(5\alpha^5 + 2\alpha^3 k^2)}{(4\alpha^4 + 5\alpha^2 k^2 + k^4)^2} \exp\left(-\frac{k^2 l' |\tau - \tau'|}{6}\right) = \frac{\alpha^3}{4\pi^{3/2}} \left( \frac{\sqrt{6}\left(6 - 5\alpha^2 l' \left|\tau - \tau'\right|\right)}{l'^{3/2} \left|\tau - \tau'\right|^{3/2}} \right.$$
$$- 8\sqrt{\pi}\alpha^3 e^{\frac{2}{3}\alpha^2 l|\tau-\tau'|} \left( \operatorname{erf}\left( \sqrt{\frac{2}{3}} \alpha \sqrt{l'} \sqrt{|\tau - \tau'|} \right) - 1 \right)$$
$$\left. - \sqrt{\pi}\alpha^3 e^{\frac{1}{6}\alpha^2 l' |\tau-\tau'|} \left( \operatorname{erf}\left( \frac{\alpha\sqrt{l'}\sqrt{|\tau-\tau'|}}{\sqrt{6}} \right) - 1 \right) \right), \quad (35)$$

which one can use to simplify the computation of equation 34. The polymer is then discretized, where the continuous contour variables $\{L, \tau_2, \tau_1, \tau', \tau\}$ are replaced with the discrete variables $\{M, m, n, p, q\}$ respectively. One can also define $x_{m,n} = l'(m,n)/l$, which is the ratio of the renormalized Kuhn length for monomers $m$ and $n$ to the actual Kuhn length $l$, therefore the equation that fully determines $x_{m,n}$ is given as

$$l^2 (m - n) x_{m,n}^2 \left( \frac{1}{l x_{m,n}} - \frac{1}{l} \right) = -\frac{\epsilon 16}{3} \pi x_{m,n}^2 \Omega_{m,n}^{\text{coh}}(M, x_{m,n}, l, \alpha) - \frac{3\sqrt{\frac{3}{2}} l^2 x_{m,n}^2 \sqrt{\frac{1}{l^5 x_{m,n}^5}}}{2\pi^{3/2}} \Omega_{m,n}^{\text{excl}}(M), \quad (36)$$

where $\Omega_{m,n}^{\text{coh}}(M, x_{m,n}, l, \alpha)$ is

$$\Omega_{m,n}^{\text{coh}}(M, x_{m,n}, l, \alpha) = \sum_{q=m+1}^{M} \left( \sum_{p=n}^{m} C(p, q, x_{m,n}, l, \alpha) \, (m - p)^2 s_q s_p \right)$$
$$+ \sum_{q=m+1}^{M} \left( \sum_{p=1}^{n-1} C(p, q, x_{m,n}, l, \alpha) \, (m - n)^2 s_q s_p \right)$$
$$+ \sum_{q=n}^{m} \left( \sum_{p=1}^{n-1} C(p, q, x_{m,n}, l, \alpha) \, (q - n)^2 s_q s_p \right)$$
$$+ \sum_{q=n+1}^{m} \left( \sum_{p=n}^{q-1} C(p, q, x_{m,n}, l, \alpha) \, (q - p)^2 s_q s_p \right), \quad (37)$$

$C(p, q, x_{m,n}, l, \alpha)$ is defined as

$$
\begin{aligned}
C(p, q, x_{m,n}, l, \alpha) = \frac{\alpha^3}{4\pi^{3/2}} \Bigg( & \frac{\sqrt{6}\,(6 - 5\alpha^2 (x_{m,n} l)\,|q-p|)}{(x_{m,n} l)^{3/2}\,|q-p|^{3/2}} \\
& - 8\sqrt{\pi}\alpha^3 e^{\frac{2}{3}\alpha^2 l |q-p|} \left( \operatorname{erf}\left( \sqrt{\frac{2}{3}}\alpha \sqrt{(x_{m,n} l)} \sqrt{|q-p|} \right) - 1 \right) \\
& - \sqrt{\pi}\alpha^3 e^{\frac{1}{6}\alpha^2 (x_{m,n} l)|q-p|} \left( \operatorname{erf}\left( \frac{\alpha \sqrt{(x_{m,n} l)} \sqrt{|q-p|}}{\sqrt{6}} \right) - 1 \right) \Bigg),
\end{aligned} \tag{38}
$$

and $\Omega^{\text{excl}}_{m,n}(M)$ is given as

$$
\begin{aligned}
\Omega^{\text{excl}}_{m,n}(M) = & \sum_{q=m+1}^{M} \left( \sum_{p=n}^{m} \frac{(m-p)^2 \omega_{q,p}}{(q-p)^{5/2}} \right) + \sum_{q=m+1}^{M} \left( \sum_{p=1}^{n-1} \frac{(m-n)^2 \omega_{q,p}}{(q-p)^{5/2}} \right) \\
& + \sum_{q=n}^{m} \left( \sum_{p=1}^{n-1} \frac{(q-n)^2 \omega_{q,p}}{(q-p)^{5/2}} \right) + \sum_{q=n+1}^{m} \left( \sum_{p=n}^{q-1} \frac{\omega_{q,p}}{\sqrt{q-p}} \right).
\end{aligned} \tag{39}
$$

### V. MATHEMATICAL DERIVATION OF $K_D$ FOR THE LATTICE BINDING MODEL

Consider a closed system volume $V$ which is discretized into $M$ voxels (small cubes) of volume $v_{\text{vox}}$, such that the centres of the voxels form a three-dimensional lattice. The cohesive units making up a patterned polymer and the cohesive units making up a patchy-particle are confined to a sub-volume $V' < V$, that is assumed, for simplicity, to be the same size for the polymer and particle and just large enough to be comparable to their overall size. These two sub-volumes are able to move around in the system volume, and when the polymer and particle volumes are not overlapping the polymer and particle do not interact, *i.e.,* only interacting when their volumes overlap.

The sub-volume is made up of $M' = MV'/V \equiv V'/v_{\text{vox}}$ voxels, which are occupied by $N_{\text{coh}}$ polymer units and $N$ particle units in the respective polymer and particle sub-volumes. For simplicitly, the polymer units are assumed uncorrelated, *i.e.,* are randomly distributed amongst the $M'$ lattice sites, whilst the particle units are assumed fixed in place (but equidistantly distributed amongst the $M'$ lattice sites). When the sub-volume containing the polymer units and the sub-volume containing the particle units overlap, individual binding events can occur through polymer and particle units occupying the same lattice site. The chance that $m < N$ polymer units become bound is weighted according to the Boltzmann factor $\exp(-\beta\epsilon' m)$, where $\beta \equiv 1/k_BT$ and $\epsilon' < 0$ (units of $k_BT$) is the binding energy between one polymer unit and one particle unit.

Thus, the partition function of the entire system can be written as

$$\begin{aligned} \mathcal{Z} &= \mathcal{Z}_0 + \mathcal{Z}_1, \\ &= \binom{M}{M'}\left(\frac{M-M'}{M'}\right)\binom{M'}{N_{\text{coh}}} + \binom{M}{M'}\sum_{n=0}^{N}\binom{M'-N}{N_{\text{coh}}-n}\binom{N}{n}\sum_{m=0}^{n}\binom{n}{m}e^{-\beta\epsilon' m}, \end{aligned} \tag{40}$$

where $\mathcal{Z}_0$ and $\mathcal{Z}_1$ are the number of possible microstates of the system when the two sub-volumes are non-overlapping and overlapping respectively and $n$ is the number of lattice sites occupied by both polymer and particle units (when the sub-volumes overlap).

Focussing on the first term (corresponding to $\mathcal{Z}_0$): the first factor is the number of ways of placing the polymer sub-volume in the system volume, the second factor is the number of ways of placing the particle sub-volume in the system volume (now containing the polymer sub-volume too), and the third factor is the number of ways of arranging the $N_{\text{coh}}$ polymer units in the polymer sub-volume (consisting of $M'$ sites).

Now focussing on the second term (corresponding to $\mathcal{Z}_1$): the first factor is the number of ways of placing the "overlapping" sub-volume (containing the polymer and particle units) in the system volume, the first binomial coefficient is the number of ways of arranging the $N_{\text{coh}}-n$ free polymer units amongst the remaining $M'-N$ sites in the sub-volume, the second binomial coefficient is the number of ways of arranging the $n$ polymer units onto the lattice sites occupied by the particle units, the second sum (over $m$) is there to account for any other requirement for the units being bound, *e.g.,* orientational/conformational alignment of the binding sites, the thirs binomial coefficient is the number of ways of arranging $m$ bound polymer units amongst the $n$ overlapping sites, and the last factor is the Boltzmann weight for a polymer unit and particle to be in the bound state. Further on, we will derive the $K_D$ expression without the sum over $m$.

Evaluating the sums in equation 40 and simplifying results in

$$\mathcal{Z} = \frac{M\left((M-M')\binom{M'}{N_{\text{coh}}} + M'\binom{M'-N}{N_{\text{coh}}}{}_2F_1\left[-N_{\text{coh}}, -N; \gamma; 1+e^{-\beta\epsilon'}\right]\right)}{M'^2}, \tag{41}$$

where ${}_2F_1[a,b;c;d]$ is the Gauss hypergeometric function, and $\gamma = 1 + M' - N - N_{\rm coh}$. The probability that at least one polymer unit is bound to a particle unit $P_1$ (defining the bound state for the system) is given as

$$P_1 = \frac{\frac{M}{M'} \sum_{n=1}^{N} \binom{N}{n} \binom{M'-N}{N_{\rm coh}-n} \sum_{m=1}^{n} \binom{n}{m} e^{-\beta\epsilon' m}}{\mathcal{Z}}, \tag{42}$$

which, when the sums are evaluated, is also expressed as

$$P_1 = \frac{M'\gamma\binom{M'-N}{N_{\rm coh}-1}\left({}_2F_1\left[-N_{\rm coh}, -N; \gamma; 1+e^{-\beta\epsilon'}\right] - {}_2F_1\left[-N_{\rm coh}, -N; \gamma; 1\right]\right)}{N_{\rm coh}\left((M-M')\binom{M'}{N_{\rm coh}} + M'\binom{M'-N}{N_{\rm coh}} {}_2F_1\left[-N_{\rm coh}, -N; \gamma; 1+e^{-\beta\epsilon'}\right]\right)}. \tag{43}$$

Therefore, the probability that no polymer units are bound (defining the unbound state for the system) is then $P_0 = 1 - P_1$. From the ratio of the unbinding and binding probabilities for the two molecules the dissociation constant $K_D$ (with units mol·nm$^{-3}$) can be written as

$$K_D = \frac{1}{N_A V}\frac{P_0}{P_1}, \tag{44}$$

where $N_A$ is Avogadro's constant. The above equation is converted to molar (M) units via 1 nm$^{-3} \equiv 10^{-24}$ liters (as in the main text for the other $K_D$s). Note that in the dilute limit approximation, where $P_0$ is close to 1 (but still less than 1), equation 44 practically gives the same result as $K_D = \frac{1}{N_A V}\frac{P_0^2}{P_1}$ suggested by the law of mass action. Plugging equation 43 into equation 44 results in

$$K_D = \frac{(M-M')\Gamma(M') + \Gamma(\gamma+N)\Gamma(\gamma+N_{\rm coh})\,\tilde{\mathcal{A}}}{N_A V \gamma \Gamma(\gamma+N_{\rm coh})\Gamma(\gamma+N)\left(\tilde{\mathcal{B}} - \tilde{\mathcal{A}}\right)}, \tag{45}$$

$$\tilde{\mathcal{A}} = {}_2\tilde{F}_1\left[-N_{\rm coh}, -N; \gamma; 1\right],$$

$$\tilde{\mathcal{B}} = {}_2\tilde{F}_1\left[-N_{\rm coh}, -N; \gamma; 1+e^{-\beta\epsilon'}\right],$$

where $\Gamma(n) = (n-1)!$ is the Gamma function and ${}_2\tilde{F}_1\left[a,b;c;d\right] = {}_2F_1\left[a,b;c;d\right]/\Gamma(c)$ is the regularized hypergeometric function.

Setting $N = 1$, *i.e.,* assuming the patchy-particle to be one cohesive unit in equation 45 results in

$$K_D = \frac{e^{\beta\epsilon'} M}{N_A N_{\text{coh}} V} = \frac{e^{\beta\epsilon'}}{N_A N_{\text{coh}} v_{\text{vox}}} = \frac{e^{\beta\epsilon'}}{N_A N_{\text{coh}}}, \tag{46}$$

where the last equality results from setting $v_{\text{vox}} = 1$ (in units of volume). Hence, the $K_D \propto 1/N_{\text{coh}}$ relationship arises.

We will now derive the result with the simplifying assumption that a polymer and particle are bound when they are overlapping, as opposed to taking into account any extra requirements for being bound (*i.e.* the sum over $m$ in equation 41). Similar to before, the partition function for the entire system can be written as

$$\begin{aligned} \mathcal{Z} &= \mathcal{Z}_0 + \mathcal{Z}_1, \\ &= \binom{M}{M'} \left(\frac{M - M'}{M'}\right) \binom{M'}{N_{\text{coh}}} + \binom{M}{M'} \sum_{n=0}^{N} \binom{M' - N}{N_{\text{coh}} - n} \binom{N}{n} e^{-\beta\epsilon' n}, \end{aligned} \tag{47}$$

where the variable $m$ is omitted. Evaluating the sums in equation 47 and simplifying results in

$$\mathcal{Z} = \frac{M\left((M - M')\binom{M'}{N_{\text{coh}}} + M'\binom{M'-N}{N_{\text{coh}}} {}_2F_1\left[-N_{\text{coh}}, -N; \gamma; e^{-\beta\epsilon'}\right]\right)}{M'^2}, \tag{48}$$

where ${}_2F_1[a, b; c; d]$ is the Gauss hypergeometric function, and $\gamma = 1 + M' - N - N_{\text{coh}}$. Equation 48 is almost identical to equation 41 except the last variable in the hypergeometric funtion is $e^{-\beta\epsilon'}$ rather than $1 + e^{-\beta\epsilon'}$. The probability for binding is then given by

$$P_1 = \frac{\frac{M}{M'} \sum_{n=1}^{N} \binom{N}{n} \binom{M'-N}{N_{\text{coh}}-n} e^{-\beta\epsilon' m}}{\mathcal{Z}} \tag{49}$$

which can then be evaluated to give

$$P_1 = \frac{M'\gamma\binom{M'-N}{N_{\text{coh}}-1} \left({}_2F_1\left[-N_{\text{coh}}, -N, \gamma, e^{-\beta\epsilon'}\right]\right)}{N_{\text{coh}}\left((M - M')\binom{M'}{N_{\text{coh}}} + M'\binom{M'-N}{N_{\text{coh}}} {}_2F_1\left[-N_{\text{coh}}, -N, \gamma, e^{-\beta\epsilon'}\right]\right)}. \tag{50}$$

As before, evaluating $K_D = (N_A V)^{-1} P_0/P_1$ (where $P_0 = 1 - P_1$) results in

$$K_D = \frac{\Gamma(\gamma+N)\Gamma(\gamma+N_{\text{coh}}) + (M-M')\Gamma(M')\Gamma(\gamma)}{N_A V \Gamma(\gamma+N)\Gamma(\gamma+N_{\text{coh}})\left({}_2F_1\left[-N_{\text{coh}}, -N, \gamma, e^{-\beta\epsilon'}\right]\right)}. \tag{51}$$

Next, setting $N = 1$ (as before) leads to

$$K_D = \frac{e^{\beta\epsilon'}(M - N_{\text{coh}})}{N_A N_{\text{coh}} V} \approx \frac{e^{\beta\epsilon'} M}{N_A N_{\text{coh}} V}, \tag{52}$$

where the last approximation is for $M >> N_{\text{coh}}$ (true for dilute limit approximation). Hence, we obtain the same relationship as in equation 46.

### VI. SUPPLEMENTARY TABLES AND FIGURES

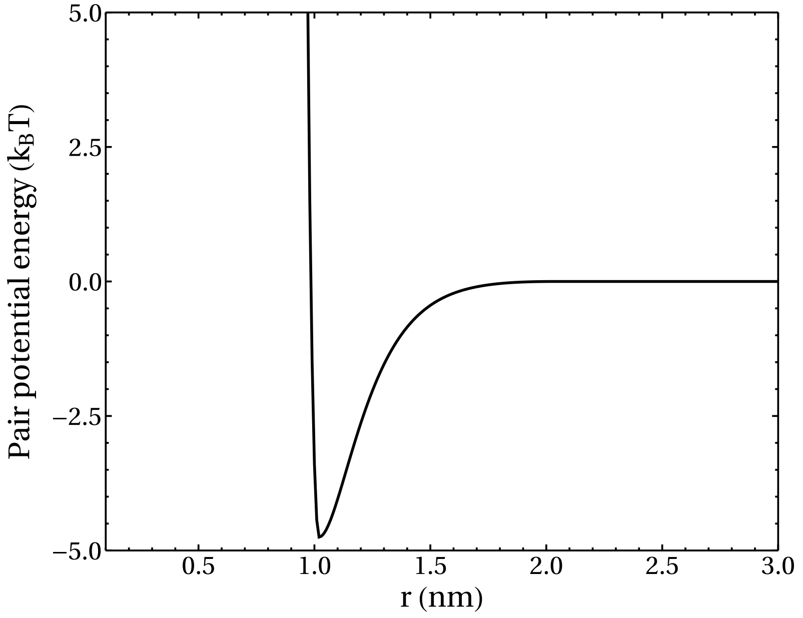


FIG. 1. Example plot of the pair potential energy as a function of distance (equation 4 in the main text). The potential shown here is for two beads at a relative distance $r$ in the 4 amino-acids-per-bead (4apb) polymer model with a well depth $\epsilon = 4.75\ k_BT$ and cut-off $2r_0 = 2 \times 1.02 = 2.04$ nm.

| $r_0$ (nm) | PCP [a] (0.38 nm) | PS [b] (3 nm) | PS (5 nm) | PB [c] (1apb) | PB (2apb) | PB (4apb) |
|---|---|---|---|---|---|---|
| PCP (0.38 nm) | 0.38 | 1.69 | 2.69 | 0.38 | 0.57 | 0.7 |
| PS (3 nm) | 1.69 | 3 | - | 1.69 | 1.88 | 2.01 |
| PS (5 nm) | 2.69 | - | 5 | 2.69 | 2.88 | 3.01 |
| PB (1apb) | 0.38 | 1.69 | 2.69 | 0.38 | - | - |
| PB (2apb) | 0.57 | 1.88 | 2.88 | - | 0.76 | - |
| PB (4apb) | 0.7 | 2.01 | 3.01 | - | - | 1.02 |

[a] PCP = Patchy-particle cohesive patch

[b] PS = Patchy-particle spacer

[c] PB = patterned polymer bead

TABLE I. Values for the distance $r_0$, used to define the excluded volume and cohesion interactions, for various combinations of beads used in MD simulations. Distances in brackets are the diameters of the respective particles. Where no simulations are performed that would include the interactions between two beads, either patchy-particle or patterned polymer, a '-' is used.

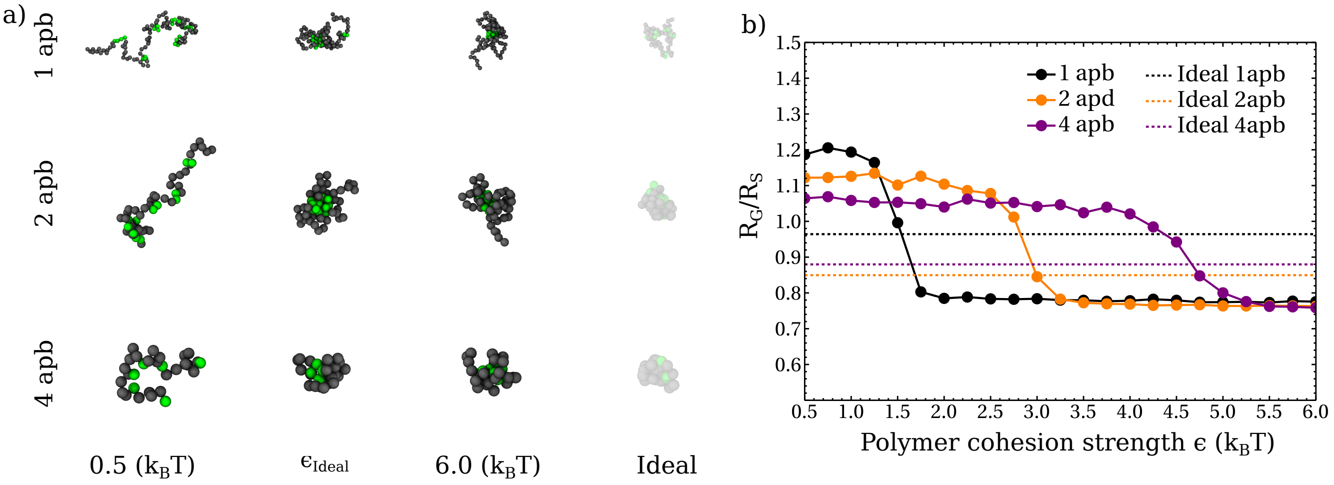


FIG. 2. Parameterization of the polymer cohesion strength $\epsilon$. **a)** MD snapshots of single patterned polymers with three choices of bead sizes (apb = amino acids per bead). **b)** Ratio of the radius of gyration to the Stokes radius as a function of polymer cohesion strength.

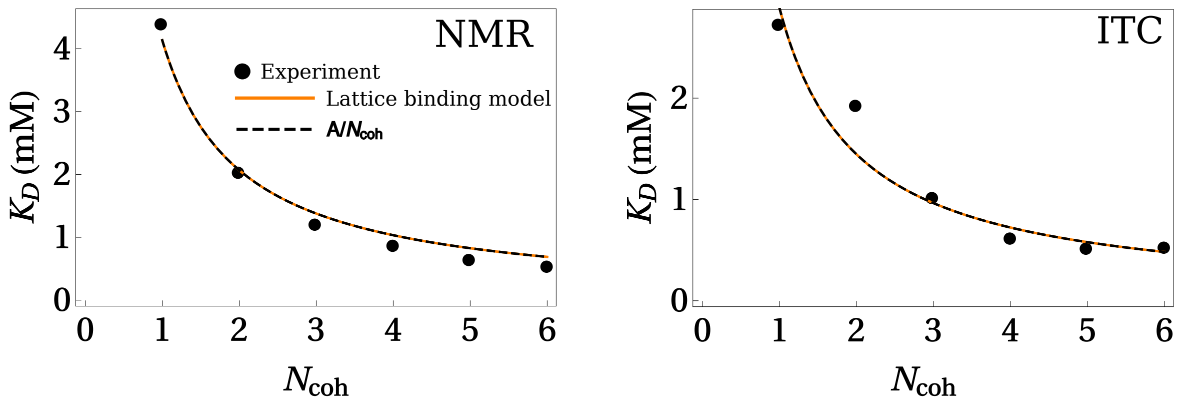


FIG. 3. Functional fits to the experimental data of synthetic FSFG sequences binding to the nuclear transport receptor NTF2 (data source [8]). (Left) Fits to the NMR data set and (right) fits to the ITC data set. The solid orange line is a fit to the data using the $K_D$ as derived in the lattice binding model (equation 52) with $N_{\mathrm{coh}}$ as given on the x-axis and $\epsilon'$ is left as a fitting parameter. The dashed line is a one parameter fit (where $A$ is a fitting constant) representing the strictly $K_D \propto 1/N_{\mathrm{coh}}$ relationship as seen in the comparison of the mean-field theory with experiments.

| Mobility parameters ($t = 10^{-2}$ μs) | NTF2 (Patchy) | NTF2 (Uniform) | Kap ($N = 4$) | Kap ($N = 10$) |
|---|---|---|---|---|
| $D$ (μm$^2$/s) | 16.4 | 12.3 | 6.9 | 5.4 |
| $D_1$ (fast) (μm$^2$/s) | 43 | 26.0 | 15.4 | 11.1 |
| $D_2$ (slow) (μm$^2$/s) | 9.2 | 7.9 | 5.4 | 4.5 |
| $w$ (slow) | 0.65 | 0.65 | 0.77 | 0.82 |
| Mobility parameters ($t = 10^{0}$ μs) | NTF2 (Patchy) | NTF2 (Uniform) | Kap ($N = 4$) | Kap ($N = 10$) |
| $D$ (μm$^2$/s) | 10.5 | 6.6 | 3.7 | 3.3 |
| $D_1$ (fast) (μm$^2$/s) | 25.7 | 21.7 | 8.1 | 6.4 |
| $D_2$ (slow) (μm$^2$/s) | 5.5 | 5.1 | 2.7 | 2.4 |
| $w$ (slow) | 0.61 | 0.82 | 0.71 | 0.68 |

TABLE II. Best fit mobility parameters for particles diffusing in a bulk solution of the F6A sequence as found using two mobility models, Gauss and BiGauss, defined by one diffusion coefficient $D$ and two diffusion coefficients $D_1$, $D_2$ respectively.

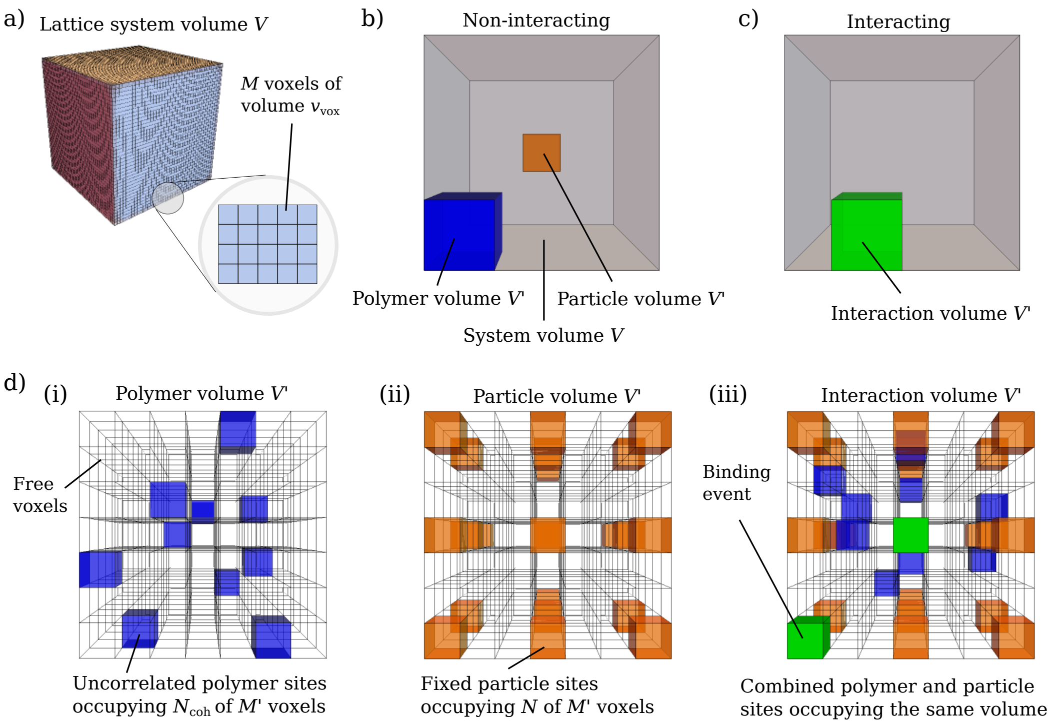


FIG. 4. Visualising the lattice binding model. **a)** System volume $V$ is discretized into $M = V/v_{\rm vox}$ voxels whose centres form a three-dimensional lattice. **b)** State of a non-interacting system. The polymer and particle are treated as sub-volumes of the system (blue and orange respectively). Since the polymer and particle volumes are not overlapping, they are not interacting. The sub-volumes are able to move freely through the volume $V$. **c)** State of an interacting system. The polymer and particle interact only when their respective volumes overlap. **d)** (i) $N_{\rm coh}$ cohesive (with respect to the particle units) polymer units can only occupy lattice sites within a sub-volume $V' < V$ (blue) which is itself free to move around the volume $V$. (ii) Particle units are fixed within a separate volume (orange) to the polymer units but of the same size $V'$. (iii) State of an interacting system. The polymer and particle sub-volumes overlap, where the uncorrelated polymer units and fixed particle units are able to interact by occupying the same voxel. The cohesive strength between a polymer and particle unit is given by $\epsilon'$.

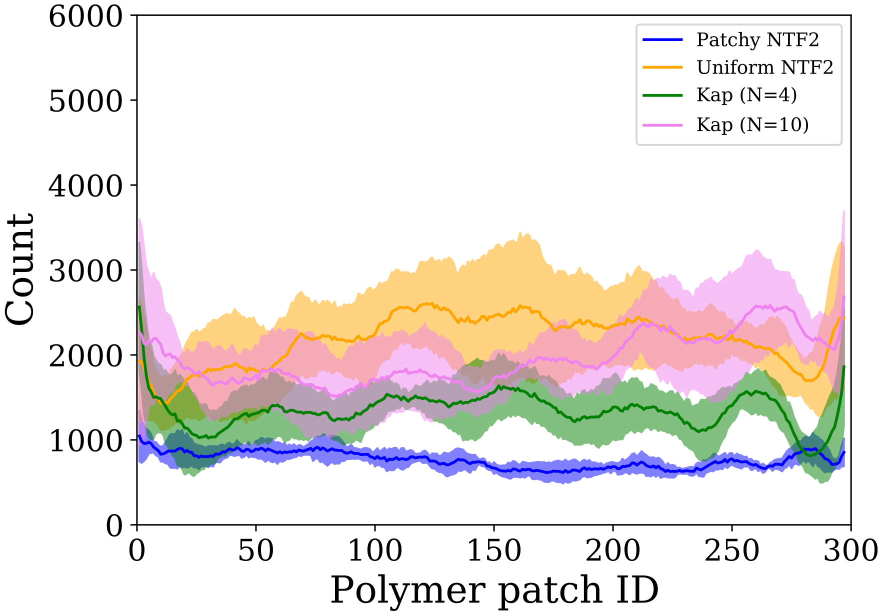


FIG. 5. Histogram data showing the number of occurences that a specific cohesive polymer bead (of the F6A sequences) is bound to a particle (within the appropriate pair potential interaction range), for a system containing one particle and many FSFG polymers at a packing fraction 0.1. If polymer were permanently stuck on the particle during the simulation, very large count peaks would be observed in the data, the relatively uniform sampling as shown here highlights that the particle is indeed binding with many polymers throughout the simulation. The smaller peaks shown for the Kap models, may be due to the particle residing for a relatively longer duration at a particular point in the simulation box. The data also shows higher counts for particles with more patches. Error bands are the standard deviation from averaging over 5 simulation runs. The polymer block ID's are ordered by polymer, where 6 polymer block IDs correspond to a single polymer (using the 4 amino acids per bead coarse-graining).

[1] Djurre H. De Jong, Lars V. Schäfer, Alex H. De Vries, Siewert J. Marrink, Herman J. C. Berendsen, and Helmut Grubmüller. Determining equilibrium constants for dimerization reactions from molecular dynamics simulations. *Journal of Computational Chemistry*, 32(9):1919–1928, 2011.

[2] Vyas Ramasubramani, Bradley D. Dice, Eric S. Harper, Matthew P. Spellings, Joshua A. Anderson, and Sharon C. Glotzer. freud: A software suite for high throughput analysis of particle simulation data, 2019.

[3] V.E. Debets, L.M.C. Janssen, and A. Šarić. Characterising the diffusion of biological nanoparticles on fluid and elastic membranes. *bioRxiv*, 2020.

[4] Alan N. Amin, Yi-Hsuan Lin, Suman Das, and Hue Sun Chan. Analytical theory for sequence-specific binary fuzzy complexes of charged intrinsically disordered proteins. *The Journal of Physical Chemistry B*, July 2020.

[5] Lucas Sawle and Kingshuk Ghosh. A theoretical method to compute sequence dependent configurational properties in charged polymers and proteins. *The Journal of Chemical Physics*, 143(8):085101, 2015.

[6] Alfredo Jost Lopez, Patrick K. Quoika, Max Linke, Gerhard Hummer, and Jürgen Köfinger. Quantifying protein–protein interactions in molecular simulations. *The Journal of Physical Chemistry B*, 124(23):4673–4685, May 2020.

[7] Harold W. Woolley. The representation of gas properties in terms of molecular clusters. *The Journal of Chemical Physics*, 21(2):236–241, February 1953.

[8] Ryo Hayama, Samuel Sparks, Lee M Hecht, Kaushik Dutta, Christina M Cabana, Jerome M Karp, Michael P. Rout, and David Cowburn. Thermodynamic characterization of the multivalent interactions underlying rapid and selective translocation through the nuclear pore complex. *Journal of Biological Chemistry*, 2018.
